# Lineage-associated tumor cell states predict outcome and guide immunotherapeutic target selection in pediatric osteosarcoma

**DOI:** 10.64898/2026.06.11.731770

**Authors:** Lindy L. Visser, Stephanie A. Schubert, Baojie Zhang, Lisa Vork Jansen, Irene F. Gosselink, Georgios Karakatsoulis, Jeff DeMartino, Laura S. Hiekcke-Jiwa, Johannes H.M. Merks, Thanasis Margaritis, Claudia Y. Janda

## Abstract

Osteosarcoma is a highly heterogeneous primary bone malignancy that predominantly affects adolescents and young adults, and robust prognostic biomarkers and effective targeted therapies remain lacking. While single-cell transcriptomics has begun to resolve tumor composition, a tumor cell–intrinsic framework linking transcriptional states to clinical outcome and therapeutic targeting is not established. Here, we present a single-cell transcriptomic analysis of pediatric osteosarcoma, integrating 22 samples across 18 patients spanning six diagnostic, six post-treatment, and ten metastatic lesions profiled using complementary sequencing platforms. Using non-negative matrix factorization, we identify recurrent lineage-associated transcriptional programs that define osteoblastic-like, chondroblastic-like, and fibroblastic-like tumor cell states. These programs show overlapping activity across tumor cells, indicating that tumor cell states are not strictly discrete. Strikingly, combined osteoblastic-like and chondroblastic-like program activity is associated with poorer overall survival, a finding validated across two independent bulk RNA-sequencing cohorts. These tumor cell states are further linked to distinct immune microenvironmental compositions. Mapping candidate immunotherapeutic targets at single-cell resolution reveals variable expression across tumor cell states, supporting state-informed strategies for CAR T cell targeting. Together, our findings establish a tumor cell state framework that links osteosarcoma heterogeneity to clinical outcome and provides a basis for patient stratification and the development of state-informed therapeutic strategies in pediatric osteosarcoma.

## Introduction

Osteosarcoma (OS) is the most common primary malignant bone tumor in adolescents and young adults, with peak incidence coinciding with the pubertal growth spurt^1,2^. The standard of care consists of neoadjuvant multi-agent chemotherapy and surgical resection, and has remained essentially unchanged for four decades. Approximately 70% of patients with localized disease achieve long-term survival, rising above 80% in patients who show good response to induction chemotherapy (histological response ≥90% tumor necrosis) and tumor-free resection margins^3,4^. However, patients with metastatic or recurrent disease face long term survival rates below 30%, and despite sustained international efforts, these outcomes have not meaningfully improved since the 1980s^3,5–7^.

A central obstacle for more effective therapies is the remarkable genomic complexity of OS. The tumor is characterized by widespread chromosomal instability, chromothripsis, and structural rearrangements. The resulting highly heterogeneous transcriptomic landscape has precluded the establishment of robust prognostic biomarkers or validated risk stratification tools as well as effective targeted therapies^8^. Histologically, conventional OS—accounting for approximately 80% of cases—is divided into osteoblastic (∼78%), chondroblastic (∼12%), and fibroblastic (∼10%) subtypes, defined by the predominant extracellular matrix produced by the malignant cells^9^. Beyond tumor-intrinsic heterogeneity, OS is embedded within a specialized bone microenvironment comprising immune, stromal, and vascular cell populations whose crosstalk with malignant cells may influence disease progression and therapeutic response^10^.

Despite these recognizable features, the molecular basis of clinical heterogeneity in OS has not been systematically characterized at single-cell resolution. Single-cell RNA sequencing (scRNA-seq) has transformed our understanding of intratumoral heterogeneity across cancer types, revealing that malignant cells organize into distinct transcriptional states—often captured as recurrent meta-programs—that carry functional and prognostic significance^11–13^. In OS, initial scRNA-seq studies have begun to resolve the cellular complexity of the tumor microenvironment^14–16^. Profiling of primary, recurrent, and metastatic lesions first revealed a tumor ecosystem composed of transcriptionally distinct malignant, stromal, and immune cell populations, with malignant cells partitioning into osteoblastic and chondroblastic populations based on canonical lineage marker expression and trajectory analyses providing evidence for transdifferentiation between these states^14^. Independent profiling of treatment-naïve tumors reinforced and refined this cellular landscape, further characterizing the immune and stromal microenvironment prior to therapeutic perturbation^15^. More recently, integration of scRNA-seq with spatial transcriptomics has extended these observations to tissue architecture, identifying spatially organized niches and candidate regions associated with chemotherapy resistance^16^. While together these studies have established insights into tumor composition and spatial organization of OS, an analysis that links recurrent malignant transcriptional states to clinical outcome and therapeutic targeting across truly independent cohorts remains lacking.

This gap is particularly consequential for patient risk stratification and the development of targeted therapies, including immunotherapy. Bulk transcriptomic studies have provided initial frameworks for prognostic classification in OS. Unsupervised analysis of tumor RNA-seq identified two prognostic groups – a favorable subgroup (G1) enriched for innate immune programs, and an unfavorable subgroup (G2) characterized by angiogenic, osteoclastic, and mesenchymal transcriptional activity – a stratification that was independently validated across multiple cohorts^17^. In parallel, tumor-intrinsic oncogenic programs, particularly MYC overexpression, have been identified as independent adverse prognostic factors, acting additively with the G1/G2 classification to define patients at higher risk^18^. Together, these findings indicate that both microenvironmental composition and malignant cell-intrinsic transcriptional programs contribute to clinical heterogeneity in OS. However, as both signatures derive from bulk tumor profiling the precise cellular origin of these prognostic signals remains unresolved, and whether they reflect distinct transcriptional states of the malignant cells themselves, or are primarily driven by the surrounding microenvironment, is unknown.

Despite extensive clinical investigation, no targeted therapy has demonstrated meaningful benefit in OS patient populations without biomarker-based selection^6,19^. The failure of these agents is unlikely to reflect universal biological resistance, but rather mismatch between therapeutic mechanism and underlying tumor biology in the absence of patient stratification. This is illustrated by the SARC028 trial, in which pembrolizumab showed a low overall response rate in OS, despite substantial variability in immune infiltration across tumors, suggesting that benefit may be restricted to molecularly defined subsets that are not captured in unselected study designs^20,21^. Similar challenges apply to CAR T cell approaches, where efficacy depends on the expression of target antigens across a heterogeneous tumor cell population^22^. Together, these observations highlight a central limitation, namely that a molecular framework linking tumor cell states to therapeutic targeting at single-cell resolution is currently lacking, limiting rational patient selection and therapy development in OS.

Here, we present a single-cell transcriptomic analysis of pediatric OS, integrating 22 samples across 18 patients, spanning pre-treatment biopsies, post-treatment resections, and metastatic lesions. Using non-negative matrix factorization (NMF), we define recurrent tumor-cell-intrinsic and lineage-associated transcriptional programs that resolve OS heterogeneity beyond discrete lineage-based classification. We show that combined osteoblastic-like and chondroblastic-like program activity is associated with clinical outcome, validated in independent bulk RNA-seq cohorts. These tumor cell states are associated with distinct immune microenvironmental compositions and varying expression of candidate CAR T cell targets. Together, these findings establish a tumor cell state framework for risk stratification and state-informed therapeutic targeting in pediatric OS, and provide a foundation for further investigation of tumor cell biology in OS.

## Results

### OS tumors display heterogeneous cellular composition with differences between primary and metastatic sites

To characterize the cellular composition of OS, we performed scRNA-seq on 22 tissues from 18 patients using two complementary platforms: SORT-seq (viably frozen samples) and 10x Genomics Gene Expression Flex (FFPE samples) (Fig. 1a). The SORT-seq cohort comprised 17 samples, including six pre-treatment biopsies, three post-treatment resections, and eight lung metastases. The 10x Flex cohort included five OS samples: three post-treatment resections, one bone metastasis, and one skin metastasis. Clinicopathological characteristics are provided in Supplementary Table S1.

**Figure 1.**
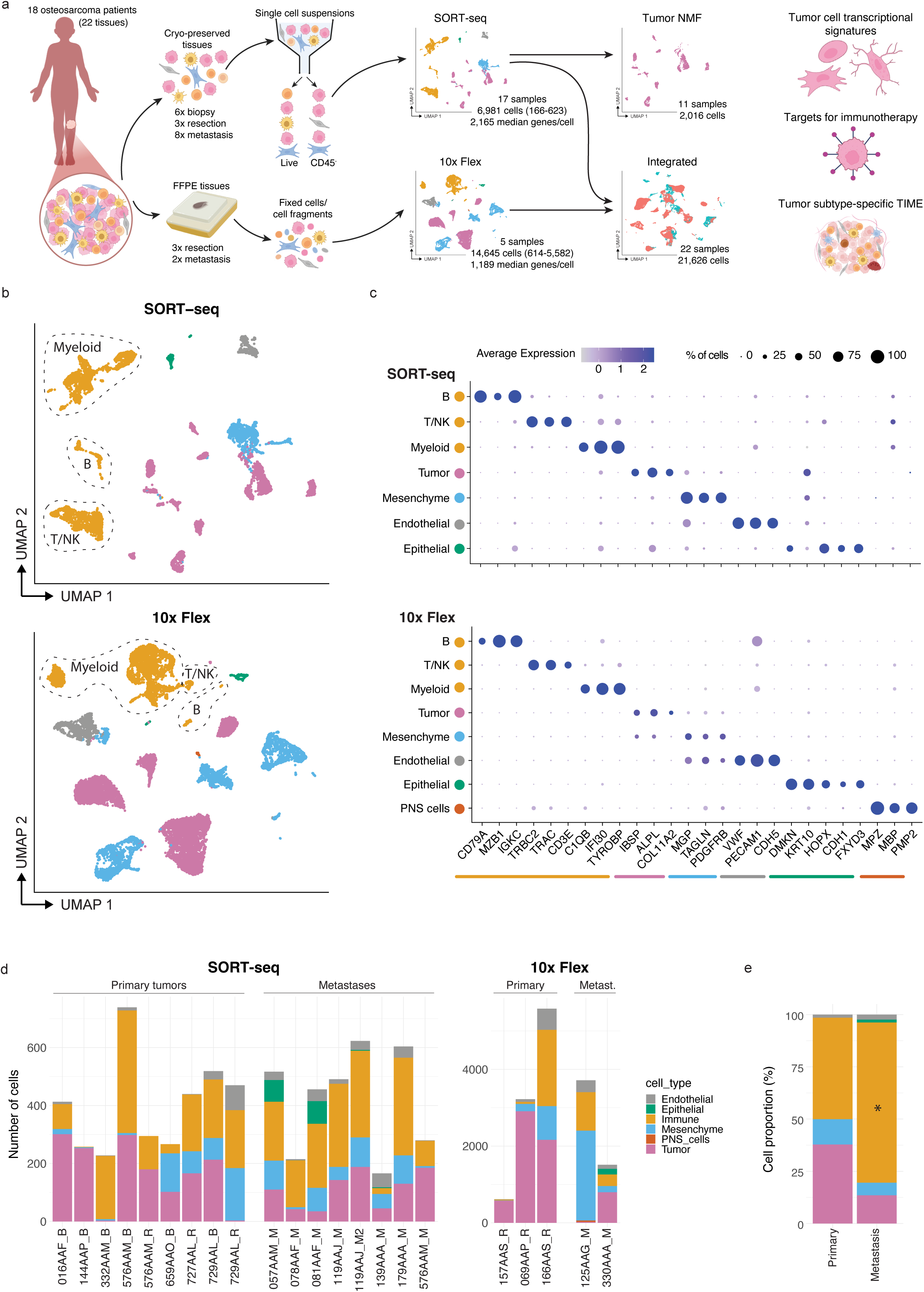
Architecture of OS tumor samples at single-cell resolution. **a.** Schematic overview of the experimental design. **b.** Uniform Manifold Approximation and Projection (UMAP) plots of the OS single-cell transcriptome from primary patient samples (top: SORT-seq dataset; bottom: 10x Flex dataset), colored by main cell type. **c.** Dot plots (top: SORT-seq dataset; bottom: 10x Flex dataset) depicting the average gene expression (dot color) of selected differentially expressed marker genes for each annotated cell type. Dot size corresponds to the percentage of cells expressing each gene. **d.** Absolute cell counts per sample (left: SORT-seq dataset; right: 10x Flex dataset), colored by main cell type. Bars are grouped by sample type. Sample labels indicate biopsy (B), resection (R), or metastasis (M). **e.** Proportion of cell types in the SORT-seq dataset per sample group. Only unbiasedly sorted samples were included. (*) indicates cell types with a significantly different proportion between primary and metastatic samples (Wilcoxon test, p<0.05).

For downstream analysis, datasets generated by the two platforms were analyzed both separately and after integration (see Methods), given the fundamental differences in their underlying chemistries. In total, scRNA-seq yielded 6,981 and 14,645 high-quality cells from the SORT-seq (166-623 cells per sample) and 10x Flex datasets (614-5,582 cells per sample), respectively. Cell clusters were annotated based on marker gene expression into six main cell types: immune cells (B cells, T/NK cells, and myeloid cells), tumor cells, mesenchymal cells, endothelial cells, epithelial cells, and peripheral nervous system (PNS) cells (Fig. 1b-c). Tumor cell identity was further confirmed by copy number variation inference (Supplementary Fig. S1), revealing distinct tumor clusters for each patient (Fig. 1b; Supplementary Fig. S2a). As expected, integration of both datasets showed substantial overlap among non-malignant cell populations (Supplementary Fig. S2a-b).

Epithelial cells were detected only in a subset of metastatic samples (Fig. 1d; Supplementary Fig. S2a) and corresponded to respiratory epithelial cells (expressing *HOPX, FXYD3*) and keratinocytes (expressing *DMKN, KRT10*) derived from lung and skin metastases, respectively (Fig. 1c). Overall tumor composition varied across samples (Fig. 1d, Supplementary Fig. S2c). However, comparison of primary tumors and lung metastases within the SORT-seq dataset revealed clear differences (Fig. 1e). Metastatic samples contained a higher proportion of immune cells (mean 77%, range 31-94%, *p*=0.036) compared to primary tumor samples (mean 49%, range 1-96%). Conversely, primary tumors tended to contain a higher fraction of tumor cells (mean 38%, range 0-98%) than metastatic samples (mean 14%, range 0-66%), although this difference did not reach statistical significance (*p*=0.11).

Due to the higher number of cells captured per sample, the 10x Flex dataset revealed additional cellular heterogeneity not observed in the SORT-seq dataset, including PNS cells (Schwann cells) and additional mesenchymal cell subpopulations (Fig. 1b-c). These included adipocytes, myocytes, mesenchymal stem cells (MSCs), osteoblasts, chondrocytes and two cancer-associated fibroblast (CAF) subsets –myofibroblastic CAFs (myoCAFs) and inflammatory CAFs– annotated based on previously defined gene signatures (Supplementary Fig. S2d-f)^16^. In contrast, mesenchymal cells in the SORT-seq dataset exhibited a more heterogeneous transcriptional profile, consistent with a mixed cell population predominantly composed of CAF-like cells (Supplementary Fig. S2e-f). Together, these data provide a comprehensive single-cell map of primary and metastatic OS, representing one of the most extensive datasets spanning diverse OS sample types and revealing pronounced inter-tumor heterogeneity.

### Lineage-associated meta-programs define distinct molecular tumor cell states in OS

Previous studies have shown that meta-programs identified across cancer types capture functionally and clinically relevant aspects of tumor biology^11,12,23^. To identify such meta-programs in the highly heterogeneous OS tumors, we analyzed transcriptional programs in tumor cells from the SORT-seq dataset (11 samples; excluding samples with fewer than 50 tumor cells and one sample with high stress signal) using NMF (Supplementary Fig. S3a; Supplementary Table S1, see Methods). This analysis identified seven consensus meta-programs (Fig. 2a). Notably, each meta-program was derived from multiple tumor samples, including both primary bone tumor and metastatic lung tissues, indicating that clustering was not driven by intra-tumor variation or sample origin/type (bone/primary versus lung/metastasis).

**Figure 2.**
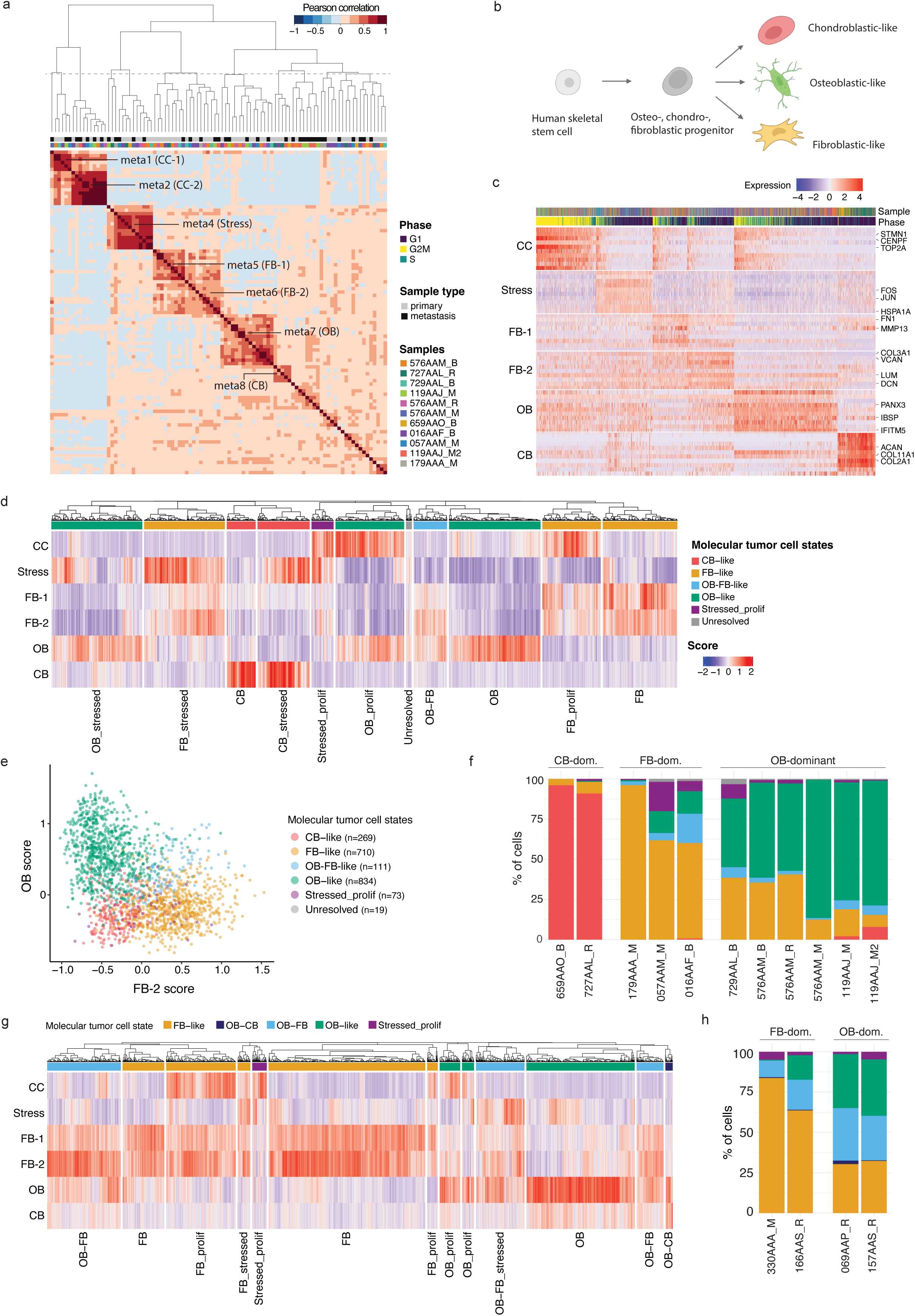
NMF defines malignant cell states in OS tumor samples. **a.** Heatmap showing the pairwise Pearson correlations between all non-negative matrix factorization (NMF)-defined transcriptional programs identified in our tumor NMF dataset (including tumor cells from 11 SORT-seq samples with >50 tumor cells). Columns are annotated by sample type and sample ID. **b.** Schematic representation of the identified molecular tumor cell types and their differentiation trajectory. **c.** Heatmap showing the scaled expression of the top 10 genes per meta-program across all cells in the tumor NMF dataset. Rows are ordered by meta-program, and columns are ordered by cell cycle phase. **d.** Heatmap depicting meta-program gene expression scores across all cells in the tumor NMF dataset. Cells (columns) are clustered based on the Pearson correlation distance of their meta-program gene expression scores. 11 clusters were identified and used to classify tumor cells into six molecular tumor cell states based on lineage (OB, CB, FB) and cell state (stressed, proliferating). **e.** Scatter plot depicting the meta-program scores of FB-2 (meta6) and OB (meta7). Points represent tumor cells, with colors indicating the assigned molecular tumor cell type based on clustering in panel d. **f.** Bar plot representing the composition of the tumor compartment per sample (NMF cohort), based on the molecular tumor cell states assigned in d. Bars are grouped by the dominant molecular tumor cell type per sample. **g.** Heatmap depicting meta-program gene expression scores across tumor cells in the 10x Flex dataset. Only samples with >50 tumor cells are included. Cells (columns) are clustered based on the Pearson correlation distance of their meta-program gene expression scores. 13 clusters were identified and used to classify the tumor cells into five molecular tumor cell states based on lineage (OB, CB, FB) and cell state (stressed, proliferating). **h.** Bar plot representing the composition of the tumor compartment per sample (10x Flex), based on the molecular tumor cell types assigned in g. Bars are grouped by the dominant molecular tumor cell type per sample. CC = cell cycle; FB = fibroblastic; OB = osteoblastic; CB = chondroblastic; OB-FB = osteoblastic/fibroblastic; OB-CB = osteoblastic/chondroblastic. Sample labels indicate biopsy (B), resection (R), or metastasis (M).

We next merged and annotated these into six largely distinct transcriptional programs (Supplementary Fig. S3b), encompassing both cell-state-related programs (cell cycle/meta1+2 and stress/meta4) and lineage-associated programs resembling non-malignant skeletal cell states (chondroblasts/meta8, osteoblasts/meta7, and fibroblasts/meta5+6; Fig. 2b, Supplementary Fig. S3c). Each meta-program defined by its top 30 weighted genes was expressed in subsets of cells across multiple tumor samples and displayed largely distinct but partially overlapping expression patterns between fibroblastic (FB-1, FB-2), osteoblastic (OB), and chondroblastic (CB) programs (Fig. 2c, Supplementary Fig. S3b). The annotation of these molecular tumor cell states was further supported by applying meta-program scoring to a single-cell mouse skeletal cell atlas^24^, with detailed annotation of normal skeletal cell types (Supplementary Fig. S3d).

We then used meta-program module scores to assign a molecular state to each tumor cell, defining three main tumor cell states: chondroblastic-like (CB-like), osteoblastic-like (OB-like), and fibroblastic-like (FB-like) (Fig. 2d-f, Supplementary Fig. S3e-f). A smaller subset of cells exhibited mixed or ambiguous lineage signatures and was therefore classified into three additional states: an osteoblastic/fibroblastic-like (OB-FB-like), reflecting the transcriptional overlap between osteoblasts and fibroblasts^25^; a stressed/proliferative state, characterized by stress-response and cell cycle gene expression; and an unresolved state, comprising cells that could not be confidently assigned to any defined lineage. Analysis of meta-program scores in the full SORT-seq dataset (malignant and non-malignant cells) revealed that OB and CB programs were enriched in tumor cells, whereas FB scores were detected in both tumor cells and non-malignant mesenchymal cells (Supplementary Fig. S4a). This observation was independently validated in the OS scRNA-seq dataset of Zhou *et al.*^14^ and also holds in our 10x Flex dataset (Supplementary Fig. S4b), supporting the robustness of OB and CB meta-programs and their suitability for analysis of bulk transcriptomic data. Projection onto our 10x Flex dataset, which contained additional non-malignant skeletal cell populations, showed the expected lineage-associated patterns, with osteoblasts displaying high OB scores, while the remaining mesenchymal populations displayed low OB scores and elevated FB-2 scores (Supplementary Fig. S4c-d). This pattern is consistent with the known transcriptional similarity between osteoblasts and fibroblast-like mesenchymal cells, which can exhibit overlapping gene expression profiles depending on differentiation states. Together, these analyses support the biological annotation of the OB, CB, and FB meta-programs as lineage-associated transcriptional programs, while highlighting that these programs can be shared between malignant and corresponding non-malignant mesenchymal populations.

Examination of tumor composition revealed substantial inter-tumor variability in the proportions of the molecular cell states (Fig. 2f). Based on the predominant cell states, tumors in the SORT-seq cohort could be classified as OB-dominant (*n*=5), CB-dominant (*n*=2), or FB-dominant (*n*=3).

The molecular cell states were independently validated in the 10x Flex dataset, where meta-program scoring identified five molecular cell states (FB-like, OB-FB-like, OB-like, stressed/proliferative and OB-CB-like tumor cells; Fig. 2g), and tumors could be classified as FB-dominant (*n*=2) or OB-dominant (*n*=2) (Fig. 2h). Together, these results define a conserved yet heterogeneous transcriptional architecture in OS, in which three lineage-associated meta-programs delineate distinct molecular tumor cell states and underlie pronounced inter-tumor variability in cellular composition.

### High OB and CB meta-program activity is associated with poorer survival in OS

To assess whether our NMF-derived molecular tumor cell states are associated with clinical outcome, we performed survival analysis. To determine which meta-program scores to include, we used a multivariable Cox proportional hazards model with stepwise variable selection, which identified OB and CB scores as the most informative risk factors. We subsequently applied the OB and CB scores to the bulk Princess Máxima Center (PMC) RNA-seq cohort, an in-house OS bulk gene expression dataset of 48 patients previously described by van Ewijk *et al.* (Supplementary Fig. S5a)^18^. Strikingly, patients with low combined OB and CB (OB+CB) scores showed improved overall survival compared to those with intermediate or high scores (*p*=0.044; Supplementary Fig. S5b). Notably, when evaluated individually, neither the OB nor the CB signatures alone was predictive of patient outcome (Supplementary Fig. S5c-d). This suggests that the combined OB+CB signal captures prognostic information that is not apparent from either program alone, despite the relatively small cohort size.

To validate the prognostic value of these meta-program signatures in an independent cohort, we analyzed the TARGET-OS RNA-seq dataset (*n*=86). Scoring patient samples based on OB and CB meta-program signatures revealed that tumors with low OB+CB scores were associated with significantly improved overall survival compared to those with high scores (*p*=0.0035; Fig. 3a-b). Together, these findings indicate that tumors enriched for OB-like and/or CB-like molecular states are associated with poorer clinical outcomes, whereas tumors with low representation of these states show more favorable survival.

**Figure 3.**
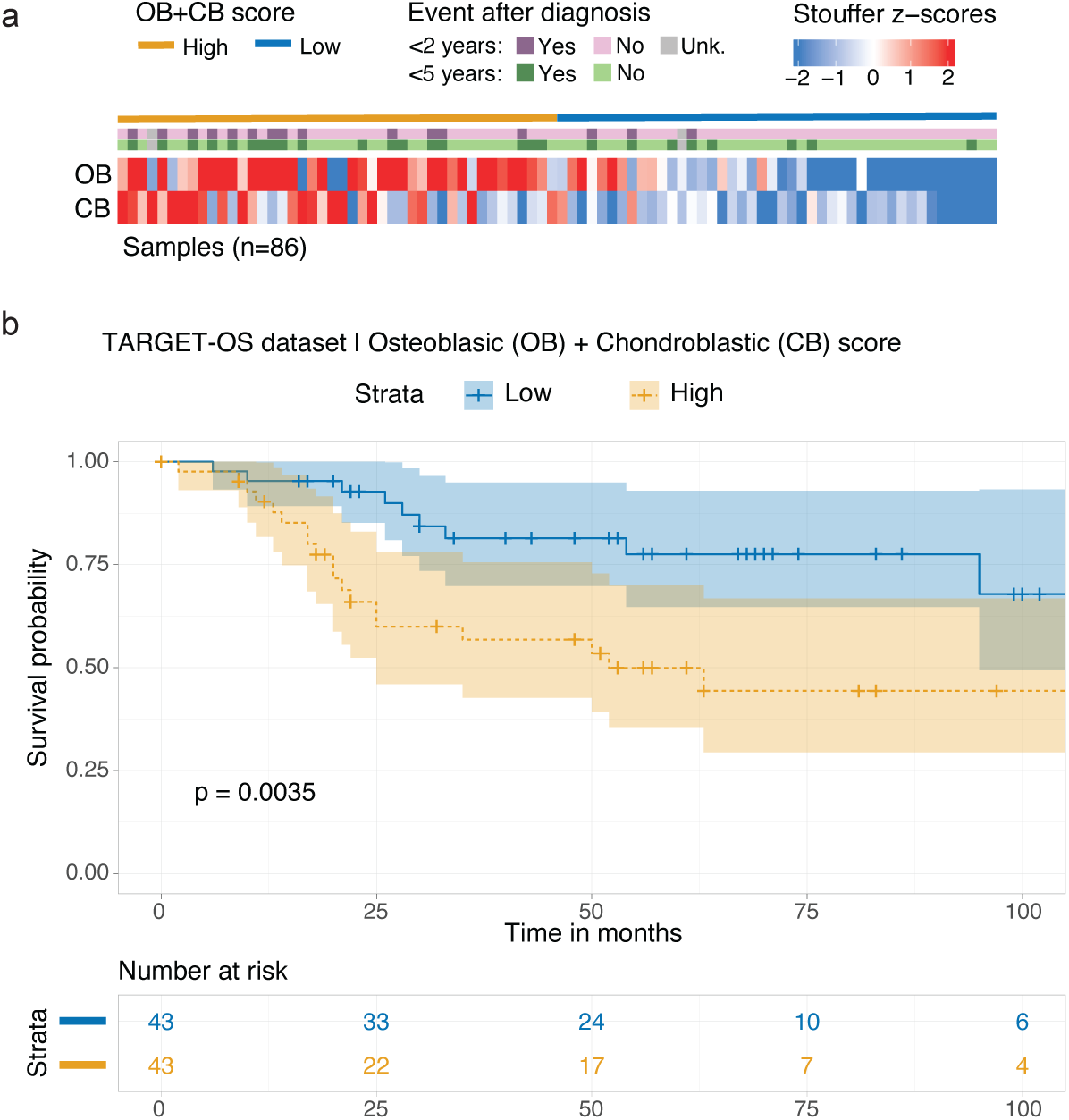
High OB and CB meta-program activity is associated with poorer clinical outcome in OS. **a.** Heatmap showing meta-program Stouffer z-scores across TARGET-OS bulk RNA-seq samples (*n=*86). Samples (columns) are ordered by combined osteoblastic and chondroblastic (OB+CB) scores from high to low. Annotation bars indicate OB+CB score groups (median split; yellow/blue), as shown in panel b, and clinical events (deaths) occurring within 2 or 5 years after diagnosis (pink and green, respectively). **b.** Kaplan-Meier survival analysis of overall survival in the TARGET-OS cohort stratified by combined OB+CB scores (median split), demonstrating patients with high OB+CB scores have poorer clinical outcome compared to those with low scores.

### IBSP immunohistochemistry captures combined OB- and CB-like tumor cell state activity

To validate the OB and CB tumor cell states at the protein level, we selected lineage-associated markers suitable for immunohistochemistry (IHC) on archived FFPE material, prioritizing genes with robust and cell type-restricted expression in our meta-program analysis (Fig. 2c, Fig. 4a, Supplementary Fig. S5e). Integrin-binding sialoprotein (IBSP, also known as bone sialoprotein), was selected as a marker expressed across both OB-like and CB-like tumor cells, while Aggrecan (ACAN) was selected as a CB-like-specific marker. IHC staining for both proteins was performed on tissue microarrays (TMAs), and protein expression was semi-quantitatively assessed using the H-score method. For each core, staining intensity was evaluated on a four-tiered scale (0=no staining, 1=weak, 2=moderate, 3=strong) and combined with the estimated proportion of positive cells (0-100%), yielding a continuous H-score ranging from 0 to 300 (Fig. 4b-c).

**Figure 4.**
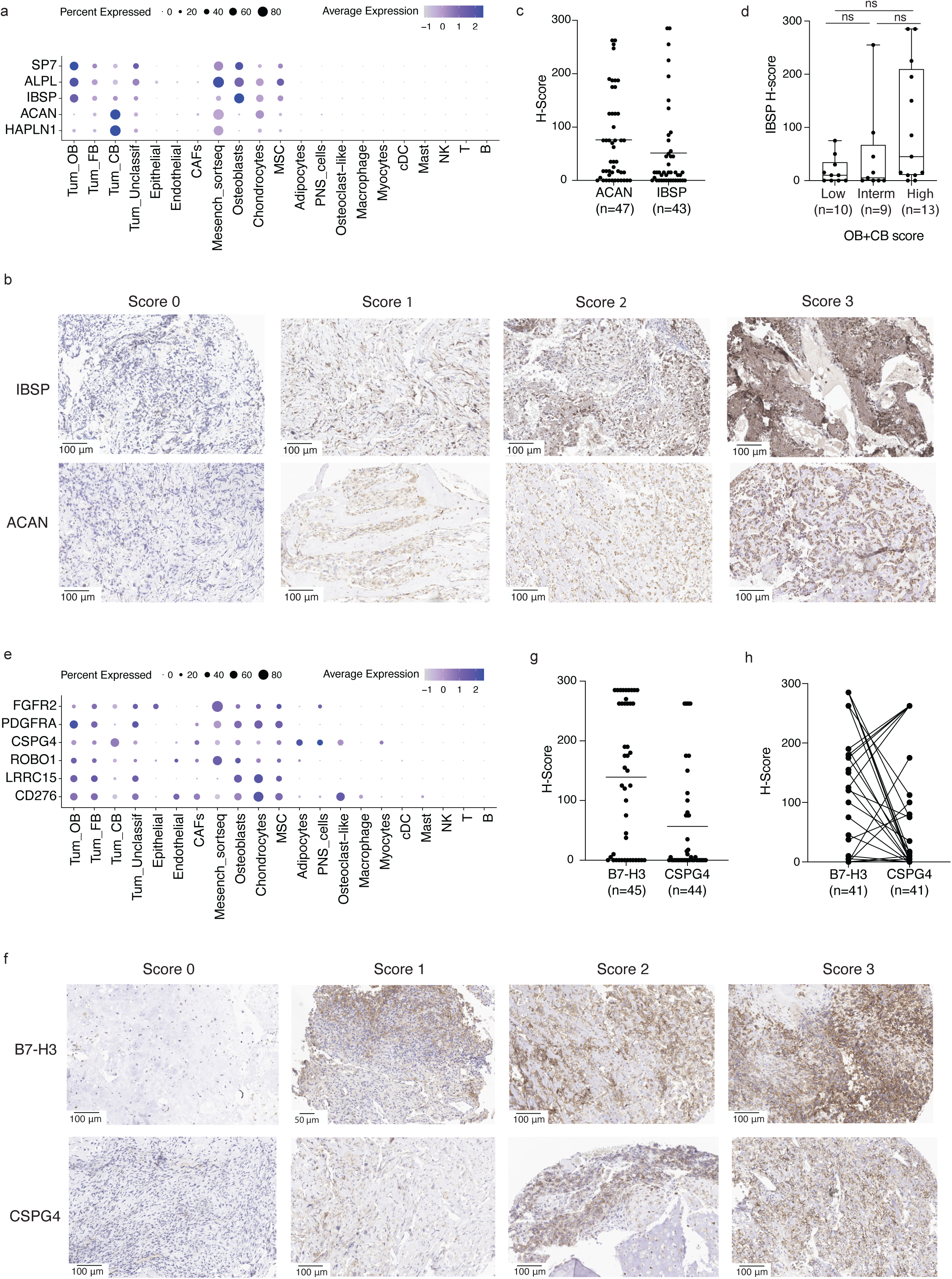
Tumor cell states guide immunotherapeutic target selection in OS. **a.** Dot plot showing average gene expression (dot color) and percentage of expressing cells (dot size) across annotated cell types for markers used for OB- and CB-like tumor cell state characterization. **b-c.** Representative IHC staining (b) and quantification (c) of IBSP and ACAN as markers for OB-like and CB-like tumor cell states in OS tissue microarrays (IBSP *n=*47, ACAN *n=*43). H-scores (range 0-300) reflect staining intensity and proportion of positive cells. **d**. IBSP H-score from TMA sections stratified by OB+CB score derived from patient matched PMC bulk RNA-seq data (low *n*=10, intermediate *n*=9, high *n*=13). Kruskal-Wallis test, *p*=0.125; Dunn’s multiple comparison, all ns. Spearman ρ=0.35, *p*=0.052. **e.** Dot plot showing average gene expression (dot color) and percentage of expressing cells (dot size) across annotated cell types for candidate immunotherapeutic targets. **f-g.** Representative IHC staining (f) and quantification (g) of B7-H3 (*n*=45) and CSPG4 (*n*=44) as candidate immunotherapeutic targets in OS tissue microarrays. **h.** Paired H-scores for B7-H3 and CSPG4 in matched tumor samples (*n*=41). Lines connect measurements from the same TMA core.

IBSP protein expression was detected in the majority of tumor samples, with H-scores ranging from 0 to 285 (Fig. 4c). To examine whether transcriptional OB+CB program activity is reflected at the protein level, OB+CB scores derived from PMC bulk RNA-seq data were correlated with IBSP H-scores from matched TMA sections in the subset of patients with both data types available (*n*=32; note these samples are distinct from the scRNA-seq discovery cohort). Stratification by OB+CB scores (low, intermediate, high) revealed a stepwise increase in IBSP H-score across score groups, with samples assigned a low OB+CB score showing a trend toward low H-scores, whereas samples with an intermediate and high OB+CB score showing a trend toward higher H-scores (Fig. 4d). Correlation analysis revealed a positive trend between IBSP H-score and OB+CB score (Spearman *ρ*=0.35, *p*=0.052, *n*=32), consistent with the expression of IBSP across both OB-like and CB-like tumor populations in our scRNA-seq data. Although this analysis will need to be extended to a larger cohort, including pre- and post-treatment samples, this concordance supports IBSP IHC as a broadly applicable tissue-based indicator of combined OB- and CB-like tumor cell state activity. In contrast, ACAN protein expression showed no meaningful correlation with the OB+CB score, consistent with its CB-restricted expression in our scRNA-seq data as well as expression of ACAN in non-malignant chondrocytes and mesenchymal cells. ACAN therefore serves as a CB-lineage-specific marker rather than a surrogate for the combined prognostic signature.

### Tumor cell states guide immunotherapeutic target selection in OS

Beyond their prognostic value for risk stratification, these molecular tumor cell states may also inform the selection of targeted therapeutic strategies, particularly for patients with poorer clinical outcome. Hence, we next explored immunotherapeutic opportunities informed by tumor cell states by examining the expression of candidate immune targets with established relevance for CAR T cell–based therapies (Fig. 4e, Supplementary Fig. S5e). Among these, the RNA of *CD276 (B7-H3)*, *LRRC15*, *ROBO1*, *CSPG4*, and *PDGFRA* were identified as broadly expressed across multiple tumor cell states, with variable expression levels and frequencies, consistent with their reported roles as tumor-associated antigens in solid cancers. Notably, while most markers showed higher expression in OB-like and FB-like tumor cells, *CSPG4* was preferentially expressed in CB-like tumor cells. To validate the expression of these targets at the protein level, IHC was performed on the two OS TMAs (Fig. 4f-g, Supplementary Fig. S5f). Unlike IBSP and ACAN, which were scored for overall protein expression, membrane restricted H-scores were applied to the CAR T cell target antigens, given that cell-surface accessibility is a prerequisite for effective CAR T cell engagement. An exception was made for ROBO1, where the selected antibody predominantly yielded cytoplasmic staining, precluding reliable membrane-specific quantification. B7-H3 showed the broadest expression with consistently high H-scores, while CSPG4 showed variable expression across cases. Consistent with their enrichment in complementary tumor cell states at the single-cell level, several tumors showed high expression of one target with low or absent expression of the other, supporting the rationale for a combinatorial B7-H3/CSPG4 targeting strategy to maximize cellular coverage across the heterogeneous OS tumor cell population (Fig. 4h). LRRC15 showed membranous protein expression in a minority of evaluated tumors, while ROBO1 was broadly expressed but predominantly restricted to cytoplasmic staining (Supplementary Fig. S5f), further supporting B7-H3 and CSPG4 as the most clinically tractable combination for cell-surface targeting in OS. Although all markers showed expression in non-malignant mesenchymal cells, B7-H3-targeting therapies have demonstrated manageable tolerability in clinical settings^26^, and CSPG4-targeting CAR T cells are currently entering first-in-human evaluation in solid tumors (NCT06096038), collectively supporting the feasibility of a combination B7-H3/CSPG4 approach in OS.

### Immune microenvironmental characteristics specific to OB- and FB-dominant samples

To assess whether the OB-, CB- and FB-dominant sample groups show differences in the immune microenvironment, the composition and functional characteristics of tumor-infiltrating immune cells were explored using graph-based clustering of the myeloid and T/NK cell compartments of the integrated OS dataset (Fig. 5a-b). Differential expression analysis and assessment of marker genes across the myeloid clusters revealed the presence of conventional dendritic cells (cDCs), mast cells, and several subsets of macrophages (Fig. 5a, Supplementary Fig. S6a). Using pro- and anti-inflammatory gene signature scoring, these macrophage subsets were predominantly skewed toward anti-inflammatory phenotypes (Supplementary Fig. S6b-d). Analysis of co-stimulatory and antigen presentation gene signatures further revealed functional differences between dendritic cell subsets, with cDC_02 exhibiting higher co-stimulatory activity and antigen presentation capacity compared to cDC_01 (Supplementary Fig. S6e).

**Figure 5.**
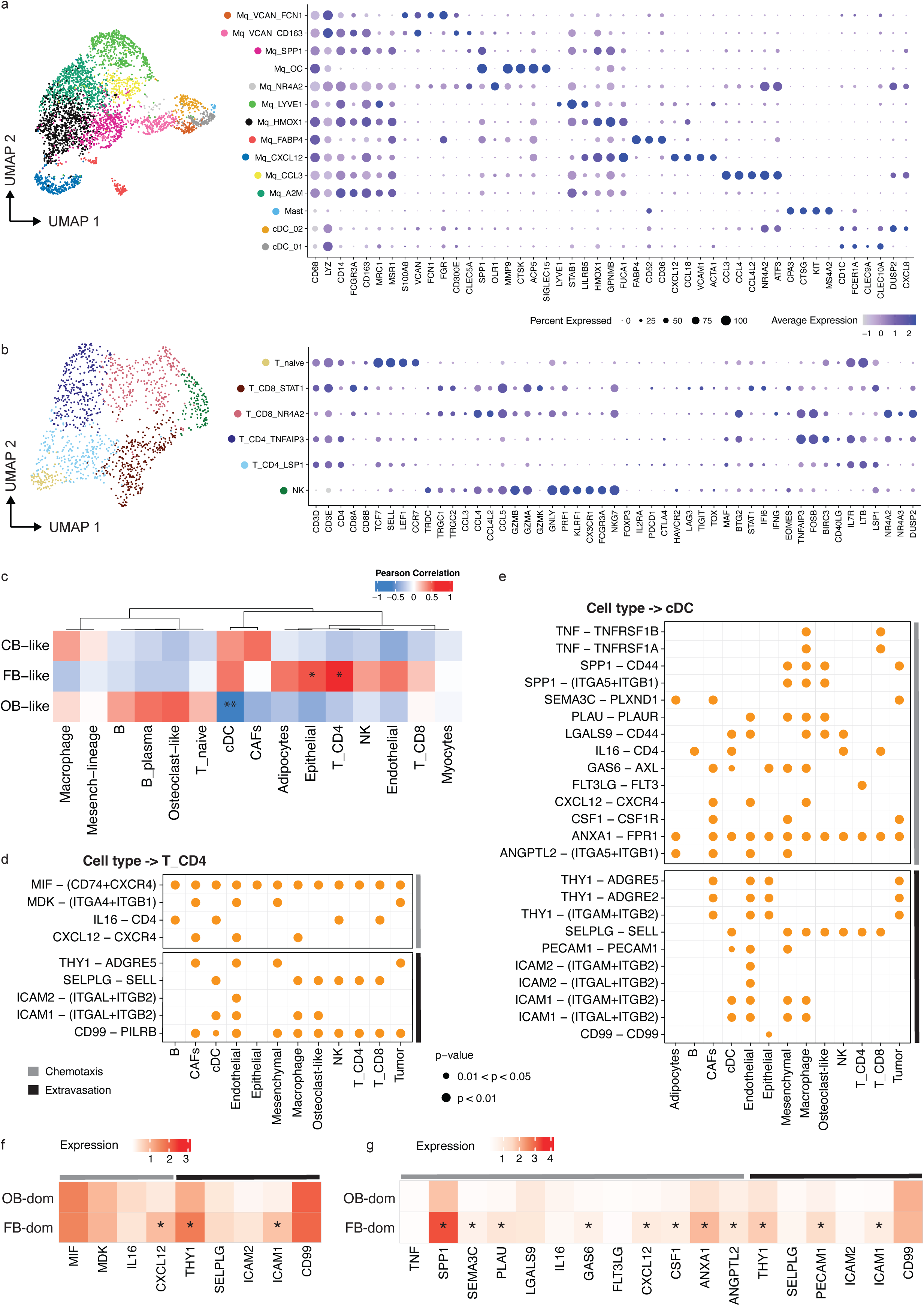
Tumor cell state-associated differences in the OS immune microenvironment. **a.** UMAP plot of myeloid cell subsets and corresponding dot plot showing average gene expression (color) of selected cell type-specific genes (Mq = macrophage; cDC = conventional dendritic cell). Dot size corresponds to the percentage of cells expressing each marker. **b.** UMAP plot of T subsets and NK cells and corresponding dot plot showing average gene expression (color) of selected cell type-specific genes. Dot size corresponds to the percentage of cells expressing each marker. **c.** Heatmap showing Pearson correlations between molecular tumor cell states and cell types within the tumor microenvironment. **d.** Bubble plot of selected ligand-receptor interactions targeting CD4+ T cells, including secreted ligands associated with chemotaxis (grey bar) and endothelial extravasation (black bar). **e.** Bubble plot of selected ligand-receptor interactions targeting cDCs, including secreted ligands associated with chemotaxis (gray bar) and endothelial extravasation (black bar). **f.** Heatmap showing the average expression of ligands involved in CD4+ T cell-targeting interactions in OB-dominant and FB-dominant samples. Genes annotated in gray correspond to chemotaxis-related interactions, whereas genes annotated in black correspond to extravasation-related interactions. **g.** Heatmap showing the average expression of ligands involved in cDC-targeting interactions in OB-dominant and FB-dominant samples. Genes annotated in gray correspond to chemotaxis-related interactions, whereas genes annotated in black correspond to extravasation-related interactions.

Among the T/NK cell clusters, several T cell subsets could be identified, including naïve T cells, CD4+ T cell subsets (TNFAIP3+ and LSP1+), and CD8+ T cell subsets (STAT1+ and NR4A2+; Fig. 5b, Supplementary Fig. S6f). Assessment of cytotoxicity and dysfunction gene signatures revealed that cytotoxic lymphocyte subsets, particularly CD8+ T cells and NK cells, exhibit high cytotoxic scores but also elevated dysfunction signatures, indicating a state of activation coupled with functional impairment consistent with a partially exhausted phenotype (Supplementary Fig. S6g).

Following identification of the immune cell subsets in the OS tumor samples, we performed correlation analysis to assess the association between molecular tumor cell states and the tumor microenvironment. FB-like tumor cell-dominant samples correlated with the presence of a higher fraction of CD4+ T cells, whereas OB-like tumor cell-rich samples were characterized by an almost complete absence of cDCs (Fig. 5c; Supplementary Fig. S6h). Epithelial cells also showed a strong correlation with FB-like tumor cells, but were not further investigated, as they were restricted to metastatic samples.

Gene set enrichment analysis of T/NK cell subsets revealed distinct immune programs associated with tumor cell states. OB+CB-dominant tumors showed enrichment of cytotoxic and activation pathways in NK and CD8+ T cells, including interferon signaling and chemokine-mediated migration, whereas FB-dominant tumors were enriched for CD4+ T cell-associated immune pathways (Supplementary Fig. S6i).

To define the mechanisms underlying these differences in immune cell composition, putative cell-cell interactions were inferred from our single-cell data using CellChat^27^. Ligand-receptor interactions between malignant cells, stratified by tumor cell state, and cells within the tumor microenvironment were modeled. This analysis highlighted several interactions enriched in FB-dominant samples that are involved in chemotaxis, such as CXCL12-CXCR4 (Fig. 5d-e, Supplementary Table S2), as well as interactions mediating immune cell extravasation through the endothelial layer, including ICAM1-ITGAL/ITGB2 (LFA-1; Fig. 5d,e; Supplementary Fig. S7a-b; Supplementary Table S2). Consistent with these findings, many of the ligands involved in chemotaxis-related interactions with cDCs and CD4+ T cells were more highly expressed in FB-dominant samples compared to OB-dominant samples, suggesting a reduced capacity of the OB-dominant tumor microenvironment to recruit these immune cell populations (Fig. 5f-g; Supplementary Fig. S7a-b). Similarly, ligands involved in extravasation-related interactions were more highly expressed in FB-dominant samples, consistent with increased expression of adhesion molecules such as ICAM1 and enhanced immune cell extravasation, whereas these signals were reduced in OB-dominant tumors.

Taken together, analysis of the OS immune microenvironment reveals both a general state of immune dysfunction, exemplified by the predominance of anti-inflammatory macrophages, and distinct tumor state–associated immune landscapes, which may in part contribute to differences in patient outcome.

## Discussion

In this study, we establish a tumor cell-intrinsic transcriptional state model in pediatric OS by integrating single-cell transcriptomic data from 22 samples across 18 patients spanning pre-treatment biopsies, post-treatment resections, and metastatic lesions, profiled using two complementary whole-transcriptome technologies. This analysis defines recurrent lineage-associated tumor cell states that resolve OS heterogeneity beyond discrete histology-based classification. We show that a combined osteoblastic (OB) and chondroblastic (CB; OB+CB) signature score is associated with clinical outcome, validated in independent bulk RNA-seq cohorts. Furthermore, these tumor cell states are linked to distinct immune microenvironmental compositions and variable expression of candidate immunotherapeutic targets. Together, these findings provide a conceptual model that links OS heterogeneity to clinical outcome and therapeutic targeting.

In previous single-cell studies of OS, including the work by Zhou *et al.*^14^, malignant cells have been classified into osteoblastic and chondroblastic populations based on canonical lineage markers, reflecting histology-like differentiation states. Within this framework, unsupervised clustering assigns cells to discrete groups, and although trajectory analyses suggest that transdifferentiation between these states can occur, the underlying classification remains categorical. Our analysis takes a complementary but conceptually distinct approach. Rather than assigning cells to fixed clusters, we apply NMF to decompose transcriptional variation into co-expressed gene programs. This approach has proven powerful for identifying recurrent meta-programs shared across tumors and linked to clinical behavior in other cancer types, including glioblastoma and melanoma^11,13^. Furthermore, Truong *et al*.^28^, previously applied NMF to define transcriptional archetypes. Yet, they relied on a mesenchymal differentiation reference derived from *in vitro* differentiated MSCs into osteoblasts, adipocytes, and chondroblasts and projected OS cells onto this normal differentiation trajectory. Therefore, this strategy does not directly capture tumor cell-intrinsic transcriptional variation across patient samples. In contrast, our approach applies NMF directly to tumor cells across a larger, clinically diverse cohort, allowing us to identify recurrent meta-programs intrinsic to the malignant compartment. Using this framework, we identified OB-like, CB-like, and FB-like states as lineage-associated transcriptional programs. Notably, OB-like and CB-like programs form more distinct transcriptional axes, whereas the FB-like program shows partial overlap with OB-like states, which may reflect shared transcriptional features between fibroblastic and osteogenic programs. This organization reveals that tumor cells are not confined to discrete lineages, but instead occupy partially overlapping transcriptional states, including intermediate OB-FB populations. The lineage-associated nature of the meta-programs we identify is independently supported by Truong *et al*.^28^, who showed that OS cells, when projected onto a normal mesenchymal differentiation atlas, express archetypes corresponding to osteogenic, chondrogenic, and mesenchymal progenitor states. Our tumor-intrinsic NMF approach converges on a similar biological organization, suggesting that these lineage programs represent a conserved feature of OS transcriptional heterogeneity.

Whether these transcriptional states represent stable cellular identities or dynamically interconvertible programs remains an open question. The observed continuum raises the possibility of tumor cell plasticity, in which cells transition between states in response to intrinsic or microenvironmental cues^13,29,30^. Addressing this will require longitudinal and functional models, such as patient-derived organoids, which enable tracking of cell state dynamics and perturbation of candidate drivers of state transitions^31^. Together, these findings provide a conceptual framework to link lineage-associated transcriptional programs to tumor behavior, clinical outcome, therapeutic vulnerabilities, and OS tumor biology.

In several pediatric cancers, molecular prognostic signatures have become integral to clinical management, enabling risk-adapted treatment that intensifies therapy for high-risk patients while reducing toxicity for those with favorable disease, for instance by MYCN-based risk stratification in neuroblastoma and FOXO1 fusion status in rhabdomyosarcoma^32–34^. OS lacks equivalent molecular tools, limiting clinical decision-making in several ways. These include, the identification of high-risk patients for enrollment in trials for novel adjuvant agents following standard therapy, the selection of low-risk patients who may be candidates for treatment de-escalation, the tailoring of post-treatment surveillance intensity based on relapse risk, and, as targeted therapies emerge, the matching of patients to treatment based on the molecular characteristics of their tumors. While several transcriptomic signatures have been proposed, the G1/G2 signature has emerged as the most clinically advanced, with independent validation across multiple cohorts and ongoing evaluation as a stratification tool in prospective trials^17,18^. The OB+CB signature described here captures a distinct and complementary dimension of tumor biology. In contrast to the G1/G2 signature, which primarily reflects microenvironmental composition, the OB+CB signature resolves malignant cell-intrinsic transcriptional states, validated in two independent bulk RNA-seq cohorts.

As the OB+CB signature is derived from meta-programs defined in malignant cells, with limited signature expression by non-malignant populations in the discovery dataset, the OB+CB signature may serve as a robust prognostic indicator across diverse clinical contexts, including diagnostic biopsies, pretreated, and relapsed tumor samples. Furthermore, the identification of IBSP as a broadly expressed OB-like and CB-like marker raises the possibility that immunohistochemical assessment could serve as a faster and more broadly accessible surrogate for transcriptomic classification in routine clinical material. The trend toward correlation between IBSP protein expression and combined OB+CB tumor cell state activity further suggests that IBSP may inform transcriptional subtype determination. Prospective validation will be required before such an approach can be adopted in practice. Together, these complementary molecular frameworks – spanning malignant cell-intrinsic transcriptional states, microenvironmental composition, and differentiation trajectory – capture distinct and non-redundant dimensions of OS heterogeneity, collectively advancing the molecular basis for patient stratification in OS.

The development of effective targeted therapies depends critically on identifying antigens preferentially expressed on the malignant cell populations driving aggressive disease^26,35–37^. The resolution of OS into molecularly defined tumor cell states now offers an analogous opportunity to evaluate whether candidate therapeutic targets are preferentially expressed in the cell populations associated with poorer outcome, and to prioritize targets accordingly. As a first step, we validated the expression of established promising clinical antigens, B7-H3^26,35^, LRRC15^38^, CSPG4^36^, and ROBO1^37^, at single-cell resolution and we found that these targets are variably expressed across tumor cell states, with CSPG4 showing preferential enrichment in CB-like cells. These findings support combinatorial targeting strategies to maximize coverage across the heterogeneous OS tumor cell population. In particular, the complementary expression patterns of B7-H3 and CSPG4 across tumor cell states — with B7-H3 preferentially expressed in OB-like and FB-like tumor cells and CSPG4 enriched in CB-like tumor cells — provide a transcriptionally grounded rationale for their combined use. Since these expression patterns are largely non-overlapping at the single-cell level, co-targeting of B7-H3 and CSPG4 would engage all three major tumor cell states and specifically address the cell populations most strongly associated with adverse clinical outcome. IHC confirmation of protein-level co-expression of both targets across the majority of evaluated OS tumors supports the feasibility of this novel combination approach. Nevertheless, several important caveats warrant consideration: relevant H-score thresholds for clinical decision-making remain to be established; the impact of decalcification on antigen preservation in bone-derived specimens may affect staining intensity; distinguishing true membranous from cytoplasmic staining can be challenging; and as only TMA cores were evaluated rather than whole-tissue sections, the extent of intratumoral heterogeneity in target expression remains to be determined. Beyond validation of known candidates, this framework of tumor cell states may also enable the systematic discovery of novel antigens enriched in tumor cells with high OB+CB program activity, providing a molecularly informed basis for next-generation target identification in OS.

In summary, this study establishes a molecular framework for pediatric OS, identifying recurrent tumor cell states associated with patient outcome, linked to distinct immune microenvironmental compositions, and informative for the prioritization of immunotherapeutic targets. These findings lay the groundwork for more precise patient risk stratification and the rational development of targeted therapies for high-risk OS patients, who continue to face dismal survival outcomes in the context of relapsed and metastatic disease.

## Materials and Methods

### Patient-derived osteosarcoma specimens

Approval for use of human material was granted by the Medical Ethical Committee of the Princess Máxima Center (Utrecht). Written informed consent was obtained from all patients and/or parents/guardians. The Máxima Biobank and Data Access Committee approved the use of tissue samples for this study (https://research.prinsesmaximacentrum.nl/en/core-facilities/biobank); project numbers: PMCLAB2018.0009, PMCLAB2025.0689, and PMCLAB2026.0791. A summary of patient demographics (age, gender), tumor characteristics (primary site, metastasis, location, outcome), and sample preparation details (platform, number of cells per sample, and analysis) is given in Supplementary Table S1.

### Processing cryopreserved tissues samples for SORT-seq

Viably frozen patient-derived tumor samples were rapidly thawed (max 60 seconds) in a water bath, minced using scalpels and mechanically disrupted by pipetting up and down, and subsequently transferred to a tube containing 5-10 ml (depending on the amount of tumor tissue) digestion buffer (Advanced DMEM/F12 [Gibco, cat no. 12634010], 20 mM HEPES buffer [Gibco, cat no. 15630056], 1 mM CaCL_2_, 100 U/mL DNase I (Stemcell, cat no. 7469) and 1 mg/mL Collagenase 1A (Sigma Aldrich, cat no. C2674; prep1) or 0.1 mg/mL Liberase DL (Sigma Aldrich, cat no. 5466202001; prep2). Samples were incubated 3x 30 min at 37°C with occasional gently swirling to expose all tissue to the digestion mixture. After each digestion cycle the sample was passed through a 70 µm filter.

After digestion, the cells were rinsed with FACS buffer (PBS + 2% FCS, 2 mM EDTA and 100 U/mL DNase I), collected in one tube and spun at 200 g for 5 min at 4°C. The cells were exposed to red blood cell lysis buffer (Invitrogen) for 5 min, rinsed, taken up in 1 mL of FACS buffer. In case non-immune cell-enrichment was required, part of the sample was stained for 30 min with human CD45 FITC-conjugated antibody (eBioscience™, cat no. 11-0459-42).

Prior to FACS sorting, single cell suspensions were stained with 5 µM DRAQ5 (Thermo Fisher, cat no. 65-0880-92) and 1 µM DAPI (Signa Aldrich cat no. D9542). CD45-negative cells viable single cells (DRAQ5-positive, DAPI-negative), were sorted, either unbiasedly or enriched for, into 384-well plates containing RT primers, using a gating strategy previously shown by Visser *et al*^39^.

### Processing of FFPE tissue samples for 10x Flex

FFPE tissue samples (1-4 years in archive) were processed according to 10x protocol: Sample Preparation from FFPE Tissue Sections for Chromium Fixed RNA Profiling (CG000632). 1-3 25 µm scrolls were used as input and were subjected to pestle dissociation. For cell counting, cells were stained with Acridine Orange/Propidium Iodide (AO/PI) and counted using a CellDrop fluorescent cell counter (DeNovix).

### Library preparation and sequencing

SORT-seq libraries were prepared according to the SORT-seq protocol^40^ and subjected to paired-end sequencing (75-bp length chemistry) using an Illumina NextSeq500 sequencer. 10x Flex libraries were prepared according to manufacturer’s protocol: Chromium Fixed RNA Profiling Reagent Kits (CG000527; 10x Genomics). Four samples were barcoded per library using Chromium Human Transcriptome Probe Set v1.0.1, with a 21-hour probe hybridization step. The 10x Flex libraries were subjected to sequencing using an Illumina NovaSeq6000 sequencer.

### Data mapping and filtering

Sequencing reads were demultiplexed and aligned to human genome GRCh38-2020-A, available from 10x Genomics (https://support.10xgenomics.com/single-cell-gene-expression/software/release-notes/build), for the SORT-seq data in R (v4.3.3) using the scruff pipeline (R package, v1.20.0), and for 10x Flex using the CellRanger pipeline (v7.2.0) from 10x Genomics^41^.

Resulting count tables were loaded in R (v 4.1.2) using the Seurat R package (v4.1.0) and were merged into one SORT-seq object and one 10x Flex object. Single cells were removed from the SORT-seq data when expressing <1,000 unique transcripts, contained mitochondrial-encoded transcripts >20% of nuclear transcripts, or >3% hemoglobin gene transcripts, and pseudogenes were removed based on GENCODE (v26) annotation. For 10x Flex, single cells were excluded if they expressed <300 unique genes, <800 unique transcripts, or contained >20% mitochondrial transcripts. DecontX was applied on both datasets to estimate and remove ambient RNA contamination in individual cells (with samples as batches), improving cross-sample comparisons^42^. Removal of cells with <1,000 and <800 unique transcripts was repeated on the decontaminated count matrix of the SORT-seq and 10x Flex data, respectively. Unique transcript counts were log normalized to 10,000 transcripts, scaled and centered. In addition, SCTransform normalization was applied for use in the dimensionality reduction. From the top 3000 most variable genes outputted by SCTransform, genes associated with cell cycle phase, sex, dissociation stress (like heat shock and chaperone), and ribosomal protein genes (gene lists provided by the SCUtils R package: https://bitbucket.org/princessmaximacenter/scutils/src/master/), as well as immunoglobin genes, were removed to avoid biases in cell clustering, as described before^43^.

The expression of the remaining top variable genes was used as input for principal component analysis (PCA) and clustering. Uniform manifold approximation and projection (UMAP) and the Louvain algorithm were used to project single-cell transcriptomes in a 2-dimensional space and determine the main cell types in the SORT-seq and 10x Flex datasets, using the first 50 principal components (PCs) and a resolution of 0.8 (Louvain algorithm) for subclustering. The cell cycle phase of each cell was inferred using the *CellCycleScoring* function of Seurat. For detailed annotation of the myeloid and T/NK compartment in the SORT-seq data, the respective clusters were subset and UMAP was rerun using 50 or 40 PCs for the myeloid and T/NK cells, respectively. To define subclusters, a resolution of 0.5 was used for the myeloid cells and 0.4 was used for T/NK cells.

### Cell type annotation

Cluster annotation was guided by SingleR, using the Human Primary Cell Atlas (^44^) reference to annotate the main cell types, and the Novershern Hematopoietic Data (^45^) and Monaco Immune Data (^46^) reference datasets to annotate the immune cell (sub)clusters. Cell type annotation was further refined by consulting cluster-specific up-regulated marker genes using the *FindAllMarkers* function of Seurat. The outputted genes were compared to known cell type-specific marker genes from literature, after which healthy cell clusters were accordingly labeled. Marker genes from Zheng *et al.*^16^ were used to annotate the cancer-associated fibroblast (CAF) clusters as myoCAFs and inflammatory CAFs.

We utilized the inferCNV package (v10.1) to infer and cluster single-cell copy number profiles (using an average expression threshold of 0.3). The immune cells, endothelial cells, CAFs and epithelial cells in our data were used as reference cells. Single-cell clusters containing CNVs were manually selected and annotated as “Tumor”.

### Data integration

The SORT-seq and 10x Flex datasets were integrated using SCT integration method from Seurat. In short, using the *SelectIntegrationFeatures* function (with parameter: reduction = “rpca”), features were selected that were variably expressed in both datasets. These features were subsequently used as “anchors” for the integration using the *FindIntegrationAnchors* and *IntegrateData* functions, and SCT normalization. The expression of the variable genes was used as input for PCA and (sub)clustering using 40 PCs and a resolution of 0.8. The subclusters annotation was guided by the cell labels of the individually analyzed SORT-seq and 10x Flex dataset and was further refined by consulting cluster-specific up-regulated marker genes from previous publications.

For detailed annotation of the myeloid and T/NK compartment, the respective clusters were subset and UMAP was rerun using 25 or 50 PCs for the myeloid and T/NK cells, respectively. To define subclusters, a resolution of 0.4 was used for the myeloid cells and 0.5 was used for T/NK cells.

### Module scoring

The Seurat function *AddModuleScore* was used to calculate module scores, considering 5 control genes per query gene.

### Non-negative matrix factorization

Non-negative matrix factorization (NMF) was performed using the RcppML R package (v0.5.6)^47^. First, tumor cells were subset from the SORT-seq dataset and samples with less than 50 tumor cells were excluded, as well as 144AAP_B since it showed a high stress signal. Then, the subsequent steps were run for every tumor sample separately.

A list of shared variable features between the individual tumor samples was compiled using the *SelectIntegrationFeatures* function from Seurat. Gene expression values were centered (but not scaled), per tumor sample, and used as input to determine the optimal NMF rank. For this the *crossValidate* function was applied, testing 2 to 20 ranks with one replicate, and was run 30 times with a different seed. The optimal rank per sample was determined by keeping the rank with the lowest error per run and taking the average of the 30 runs per sample returning one value. NMF was then run (*nmf* function), using the sample-specific optimal ranks, with a maximum of 5000 iterations, and run 50 times with different seeds. The *mse* function was utilized to return the mean squared error per run, from which for each sample the NMF run with the lowest error was selected as final NMF output.

The NMF weighted genes were subset from the output for all samples and merged, all missing values were set to zero, and gene weights were scaled (but not centered) per NMF program. Pairwise Pearson correlation coefficients were calculated between NMF-defined transcriptional programs (across all tumors), which subsequently were hierarchically clustered to determine groups (as shown in Fig. 2a). Highly correlated groups were merged into meta-programs by combining the gene lists and averaging the gene weights. Cell type scoring was performed using the top 30 weighted genes per meta-program to calculate module scores. Cell state classification was guided by hierarchical clustering of single-cell meta-program scores using a Pearson distance method (as shown in Fig. 2d,g) and plotted using the ComplexHeatmap R package.

### Gene list enrichment analysis

Functional enrichment analysis was performed on NMF-derived gene lists with the *compareClusters* function from the clusterProfiler R package (v4.2.2). The top 5 signaling pathways were plotted in bar graphs using the ggplot2 R packages (v3.3.5), showing the -log_10_ adjusted *p*-values.

### Immunohistochemistry staining

Three micrometer thick sections of FFPE blocks were cut, mounted on precoated slides, and dried for at least 30 min at 65°C. An automated BOND-RX system (Leica Microsystems) was used for the deparaffinization and IHC staining of biomarkers CSPG4, LRRC15, ACAN, IBSP, B7H3, and ROBO1. Antigen retrieval was performed by boiling the sections in Tris/EDTA (BOND Epitope Retrieval Solution 2, pH9; Leica Biosystems) for 5min for ACAN, 15 min for ROBO1 and 20 min for CSPG4 and B7H3. Antigen retrieval for LRRC15 and IBSP was performed by boiling the sections in a citrate-based buffer (BOND Epitope Retrieval Solution 1, pH 6; Leica Biosystems) for 30 min. The following primary antibodies were used: ROBO1 (clone 10E2-R, Invitrogen Antibodies, 1:200, retrieval buffer Tris/EDTA pH9); CSPG4 (clone EPR9195, Abcam, 1:400, retrieval buffer Tris/EDTA pH9); LRRC15 (clone E4X8J, Cell Signalling, 1:40, retrieval buffer citrate pH6); B7H3 (clone D9M2L, Cell Signalling, 1:40, retrieval buffer Tris/EDTA pH9); ACAN (clone 6B4, Abcam, 1:1000, retrieval buffer Tris/EDTA pH9); IBSP (polyclonal, LifeSpan BioSciences, 1:600, retrieval buffer citrate pH6). BondTM Primary Antibody Diluent (Leica) was used to dilute the primary antibodies. The sections were incubated at room temperature for 5 min for ACAN, 20 min IBSP, 30 min for CSPG4, ROBO1 and B7H3 and 60 min for LRRC15. The sections were then incubated for 8 min with a post-primary rabbit anti-mouse linker followed by incubation for 8 min with anti-rabbit horseradish peroxidase-labeled polymer. Endogenous peroxidase was blocked with 0.3 % hydrogen peroxide in deionized water for 5 min, followed by incubation for 10 min with diaminobenzidine (DAB) and again 5 min with DAB enhancer. After each incubation step, ACAN was flushed twice for 5 min using Tris Buffered Saline with Tween 20 (TBST-1X). This was done three times for ROBO1 after every incubation step, to decrease background staining. All sections were counterstained with hematoxylin (BOND Polymer Refine Detection Kit; Leica Biosystems) for 5 min, dehydrated, cleared and mounted. A positive and negative control tissue was included in each section. Under the Leica DM 2500 light microscope (Leica Co), the stained sections were scored according to the four-point scale based on the IHC intensity. Results are summarized in Supplementary Data.

### Ligand-receptor interactions analysis

The CellChat (v2) algorithm was used to model in an unbiased manner cell-cell interactions per molecular cell type-dominant group^27^. Extravasation-related interactions were subset from the output data by filtering for interactions with cDC or T_CD4 as target cell group. The resulting interactions list was further short-listed using literature. For chemokine-related interactions the output was additionally filtered for interactions annotated as “Secreted Signaling” and further short-listed using literature.

### Comparison with data from Zhou *et al*

10x single-cell RNA-seq data from the publication of Zhou *et al.*^14^, was downloaded from the NCBI Gene Expression Omnibus database (accession code GSE152048) and loaded into Seurat. We applied the QC filtering as specified in the paper (excluding cells expression <300 unique genes or >10% mitochondrial transcripts), and applied scDblFinder (v1.15.2) to remove doublets. Subsequent data normalization and variable feature filtering was performed as described for the data presented in this study. Main cell type annotation was inferred using SingleR, and meta-program scores were applied on the full dataset.

### Survival analysis

To determine which meta-program scores to include in the survival analysis, a multivariable cox proportional hazard model with stepwise variable selection (based on Akaike Information Criterion, AIC) was performed on the single-cell data. For this, the *AggregateExpression* function from Seurat was used to sum gene counts of all cells for each patient. Using this pseudobulk data we calculated the meta-program scores per patients. Individuals that did not experience the event were censored at their last follow-up time. The model pointed to a final model including OB and CB scores to have most predictive power.

The TARGET Osteosarcoma (TARGET-OS) dataset was downloaded and loaded into R using the TCGAbiolinks R package (v2.37.1). Count data files were merged into one. Mitochondrial genes, as well as genes present in <2 samples with <5 reads and with a total count (over all samples) below 20 reads, were excluded. Counts were log2 normalized using the DESeq2 R package (v1.34.0). Then, the normalized count tables were subset to contain only the genes included in the NMF-derived OB and CB meta-program signatures. Mean gene expression levels were calculated per patient and the expression values were scaled and centered per gene (i.e. *Z*-scored). Each set of 30 gene meta-program *Z*-scores was combined using Stouffer’s *Z*-score method into a signature score. Heatmaps were generated using the ComplexHeatmap R package (v2.11.1). The same analysis steps were taken for the PMC osteosarcoma bulk RNAseq dataset.

Survival analysis was performed using the survival R package (v3.2-13) and p-values were calculated using the Log-Rank test. Kaplan-Meier plots were generated using the survminer R package (v0.4.9). For the analysis, a significance level of 5% was used.

## Data availability

The scRNA-seq data of the osteosarcoma cryopreserved (SORT-seq) and FFPE (10x Flex) tissue samples, generated for the purpose of this study, will be made available upon publication. The publicly available scRNA-seq data of osteosarcoma tumors generated by Zhou *et al.*^14^ and used in this study are deposited in the NCBI Gene Expression Omnibus database under accession number GSE152048. The publicly available osteosarcoma bulk RNA-seq cohort of the Princess Máxima Center for Pediatric Oncology used in this study is deposited in the European Genome-phenome Archive (EGA) under accession number EGAS00001008073. The publicly available TARGET-OS dataset is available via open access at https://ocg.cancer.gov/programs/target/projects/osteosarcoma.

## Code availability

All analysis scripts will be made available upon publication.

## Acknowledgements

We are profoundly grateful to the patients and parents who agreed to participate in our research. The work was supported by Stichting Loeka (to C.Y.J.), the foundation of the Princess Máxima Center (to C.Y.J.), a collaborative grant initiative between the Princess Máxima Center, Utrecht University, and the University Medical Center Utrecht, as well as PPP Allowance to the Miltenyi Máxima Immune Oncology / MIMO project (LSHM24013 to C.Y.J) made available by Health-Holland, Top Sector Life Sciences & Health to stimulate public-private partnerships. We want to thank the Single Cell Genomics Facility and Prof. Dr. Frank Holstege for thoughtful discussions and helpful feedback, the Big Data Core and Dr. Roelof van Ewijk for providing RNA-seq and patient-linked survival data, and the Pathology Department and the Flow Cytometry and Cell Sorting Core Facilities for their assistance with tumor tissue staining and FACS experiments.

## Declaration of the use of Gen AI

In the final steps of the preparation of this manuscript the authors used Grammarly, Claude, and ChatGPT to improve the language. After using these tools, the authors reviewed and edited the content as needed and take full responsibility for the content.

## Author contributions

C.Y.J., T.M., and L.V. conceived the project; C.Y.J., T.M., and J.H.M.M. secured funding; C.Y.J., T.M., and J.H.M.M. supervised the research. L.S.H., L.V.J., and I.F.G. prepared samples for scRNA-seq for the SORT-seq platform, and L.S.H. and B.Z. for the 10x Flex platform. L.V. analyzed scRNA-seq data and interpreted results, with help of G.K., J.D., and S.A.S. L.V., S.A.S. and C.Y.J. wrote the manuscript, with input from all authors. All authors reviewed and approved the final manuscript.

## Supplementary figure legends

**Figure S1.**
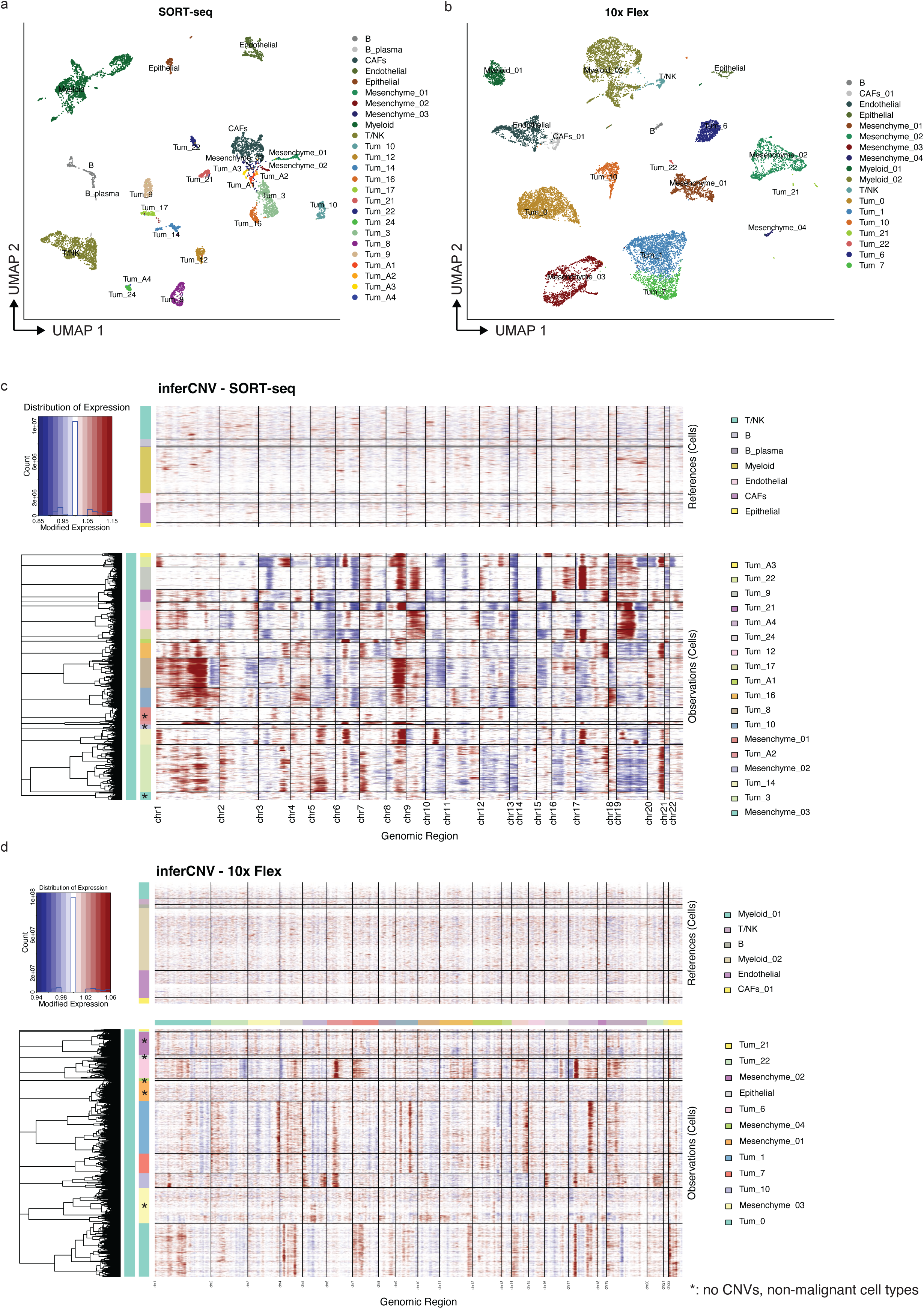
Tumor cell annotation based on inferred copy-number profiling. **a-b.** UMAP plots showing the identified clusters of the SORT-seq dataset (a; *n*=25 clusters) and the 10 Flex dataset (b; *n*=18 clusters). **c-d.** Heatmap depicting inferred copy-number variation (CNV) profiles per cell using inferCNV (c: SORT-seq dataset; d: 10x Flex dataset). Chromosomes are arranged along the horizontal axis and cell clusters along the vertical axis. Cell types indicated in the upper panels were used as reference non-malignant populations. Clusters marked with an asterisk in the lower panels were annotated as non-malignant based on the absence of CNV patterns.

**Figure S2.**
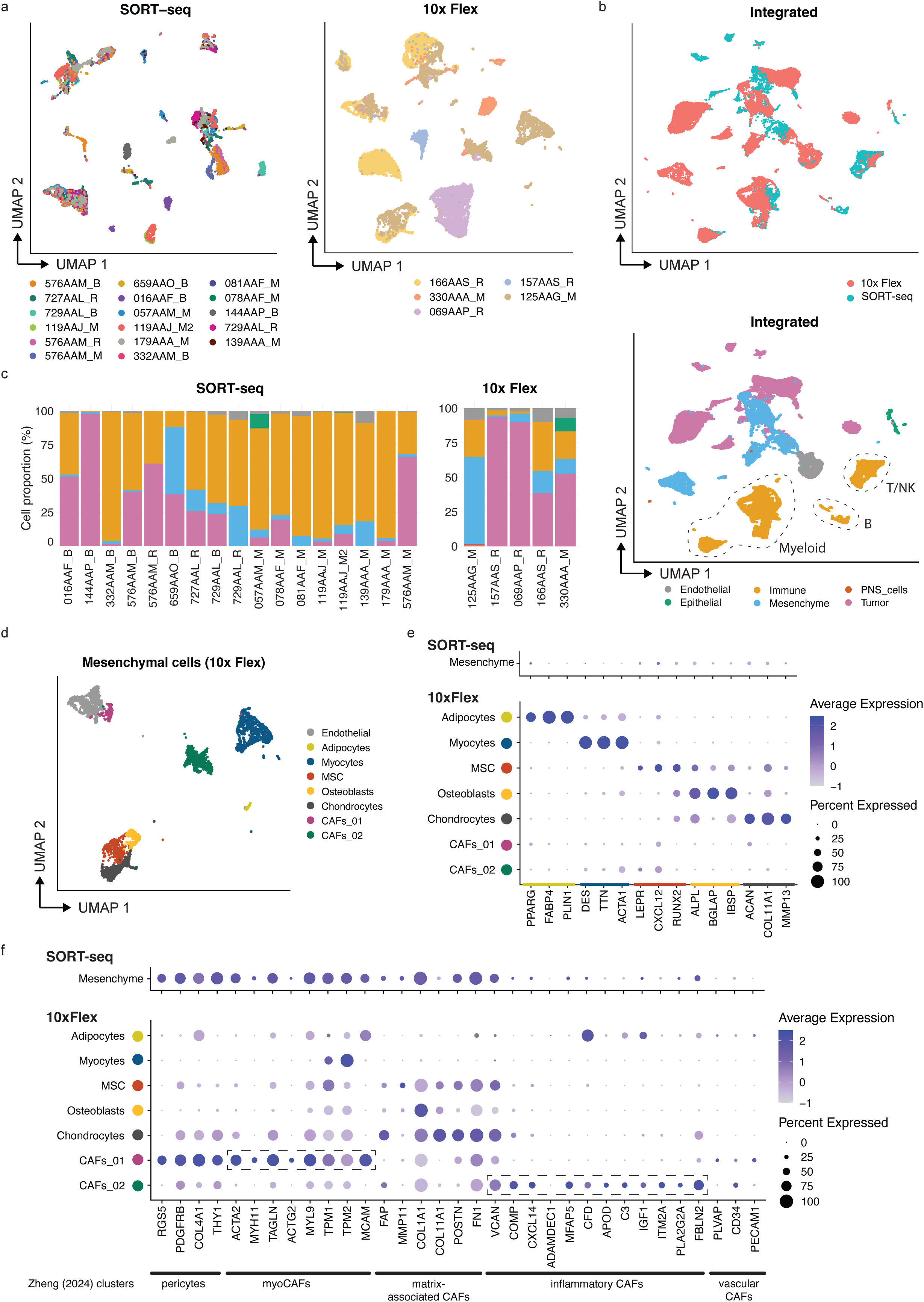
Integration and characterization of cellular heterogeneity across OS samples. **a.** UMAP plots colored by sample origin (left: SORT-seq dataset; right: 10x Flex dataset). **b.** UMAP plot of the integrated SORT-seq and 10x Flex dataset, colored by dataset of origin or main cell type. **c.** Proportion of cell types per sample (left: SORT-seq; right: 10x Flex dataset), colored by main cell type and grouped by sample type. For the SORT-seq samples, only unbiasedly sorted samples were included. **d.** UMAP plot highlighting endothelial and mesenchymal cell subsets identified in the 10x Flex dataset. **e.** Dot plots of mesenchymal cell subsets (top: SORT-seq dataset; bottom: 10x Flex dataset) showing average expression (dot color) of selected marker genes for adipocytes, myocytes, mesenchymal stem cells (MSC), osteoblasts, and chondrocytes. Dot size indicates the percentage of cells expressing each gene. **f.** Dot plots of the mesenchymal cell subsets (top: SORT-seq dataset; bottom: 10x Flex dataset) showing average expression (dot color) of mesenchymal lineage markers from Zheng, *et al*. (2024)^48^. Sample labels indicate biopsy (B), resection (R), or metastasis (M).

**Figure S3.**
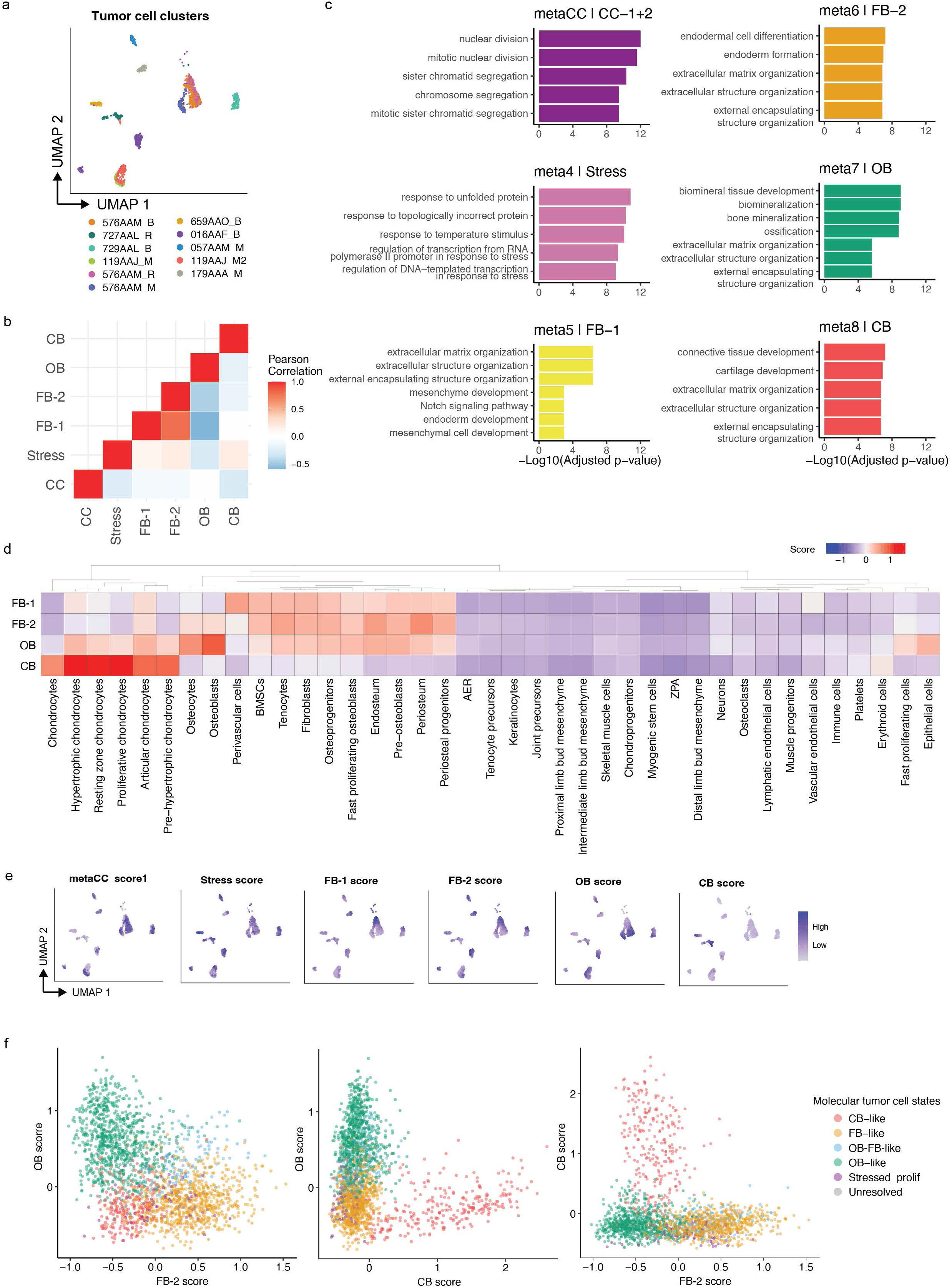
Identification and validation of transcriptional meta-programs in OS tumor cells. **a.** UMAP plot showing clusters of the tumor NMF cohort, colored by sample ID. **b.** Heatmap showing the pairwise Pearson correlations between single-tumor cell meta-program scores. **c.** Bar plots showing pathway enrichment results of the top 30 genes per meta-program. The top five pathways per meta-program are shown with corresponding -log10 adjusted *p-*values (two-sided Fisher’s exact test with Benjamini-Hochberg correction), colored by meta-program; **d.** Meta-program gene expression scores projected onto a mouse skeletal cell atlas^24^. **e.** UMAP plots showing the meta-program scores across clusters of the tumor NMF cohort. **f.** Scatter plot depicting meta-program scores of FB-2 (meta6), OB (meta7), and CB (meta8). Each point represents a tumor cell, colors according to its assigned molecular tumor cell type on clustering in Fig 2d. This panel represents an extended version of Fig 2e. Sample labels indicate biopsy (B), resection (R), or metastasis (M).

**Figure S4.**
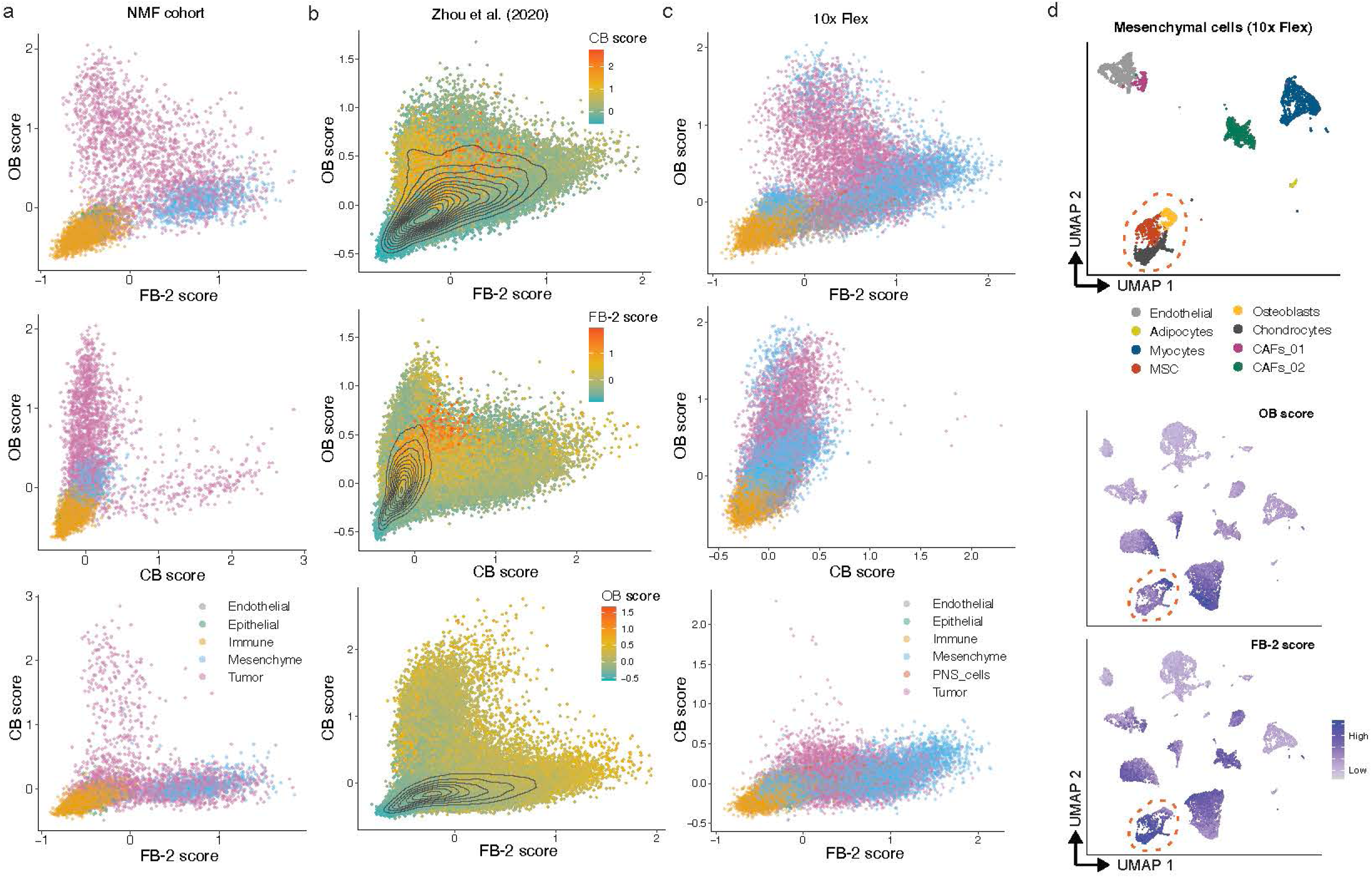
OB, CB and FB meta-program activity across tumor and mesenchymal cells. **a-c.** Scatterplots showing FB-2, OB, and CB meta-program scores across cells from the (a) NMF/SORT-seq dataset, (b) Zhou, *et al*. 2020 dataset^14^, and (c) 10x Flex dataset. Each point represents a single cell (including all cell within each dataset) with colors in panel a and c indicating the main cell types as defined in Fig. 1b. **d.** UMAP plots of the mesenchymal cells from the 10x Flex dataset (top), alongside OB and FB- 2 meta-program scores (middle and bottom), highlighting clusters corresponding to osteoblasts, mesenchymal stromal cells (MSCs), and chondrocytes.

**Figure S5.**
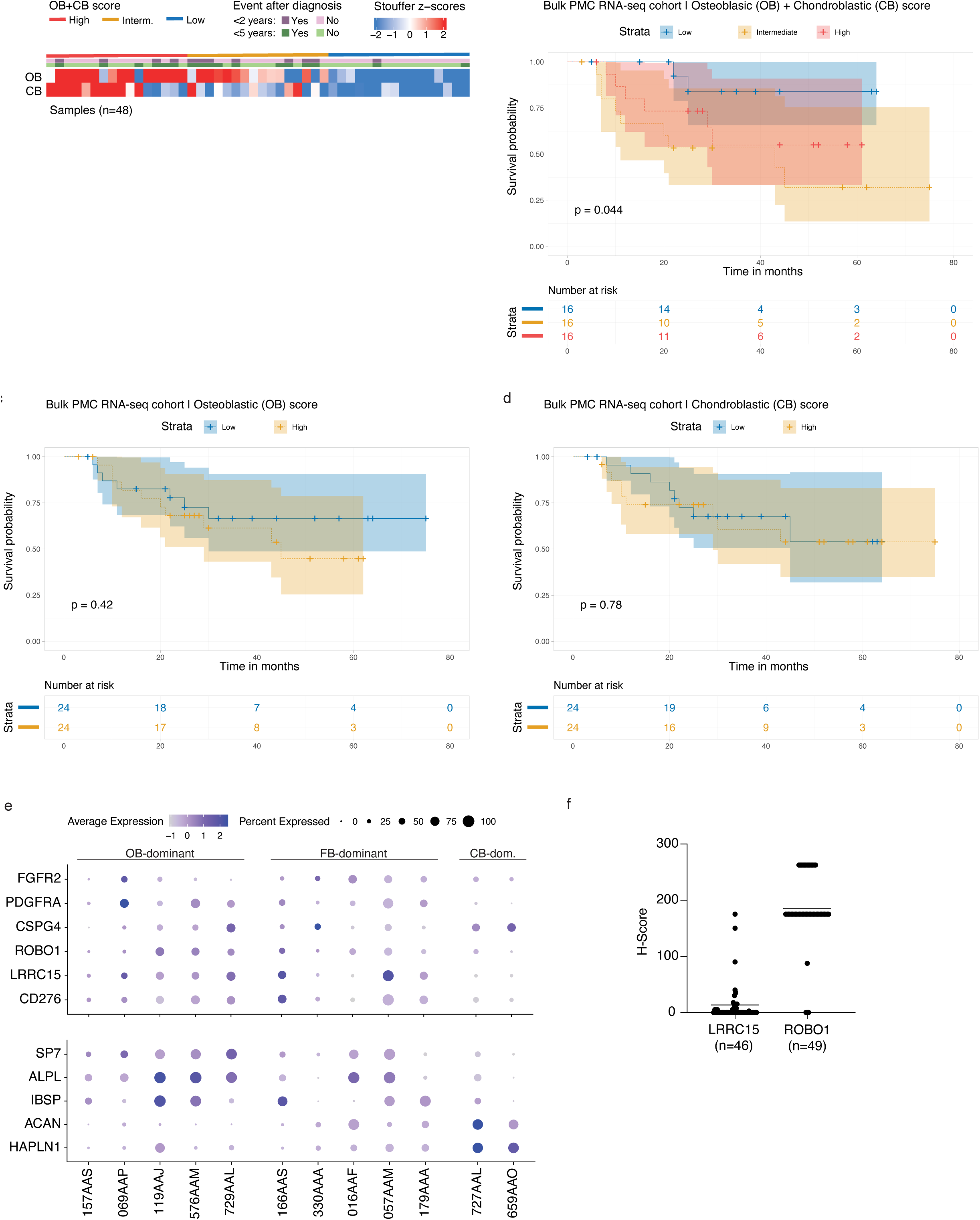
Combined OB and CB meta-program scores predict clinical outcome. **a.** Heatmap showing meta-program Stouffer z-scores across PMC bulk RNA-seq samples (*n*=48). Samples (columns) are ordered by combined osteoblastic and chondroblastic (OB+CB) scores from high to low. Annotation bars indicate OB+CB score groups (median split; red/yellow/blue) as shown in panel b, and clinical events (deaths) occurring within 2 or 5 years after diagnosis (pink and green, respectively). **b.** Kaplan-Meier analysis of overall survival stratified by combined OB+CB scores, showing separation between score groups. **c.** Kaplan-Meier analysis of overall survival based on the OB score (osteoblastic/meta7) alone in the PMC bulk RNA-seq dataset (*n*=48). **d.** Kaplan-Meier analysis of overall survival based on the CB score (chondroblastic/meta8) alone in the PMC bulk RNA-seq dataset (*n*=48). **e.** Dot plot showing the average gene expression (dot color) across tumor cells per individual for (top) markers of molecular tumor cell types and (bottom) candidate therapeutic targets. Dot size indicates the percentage of tumor cells expressing each gene. **f.** Quantification of IHC staining of LRRC15 (*n*=46) and ROBO1 (*n*=49) as candidate immunotherapeutic targets in OS tissue microarrays.

**Figure S6.**
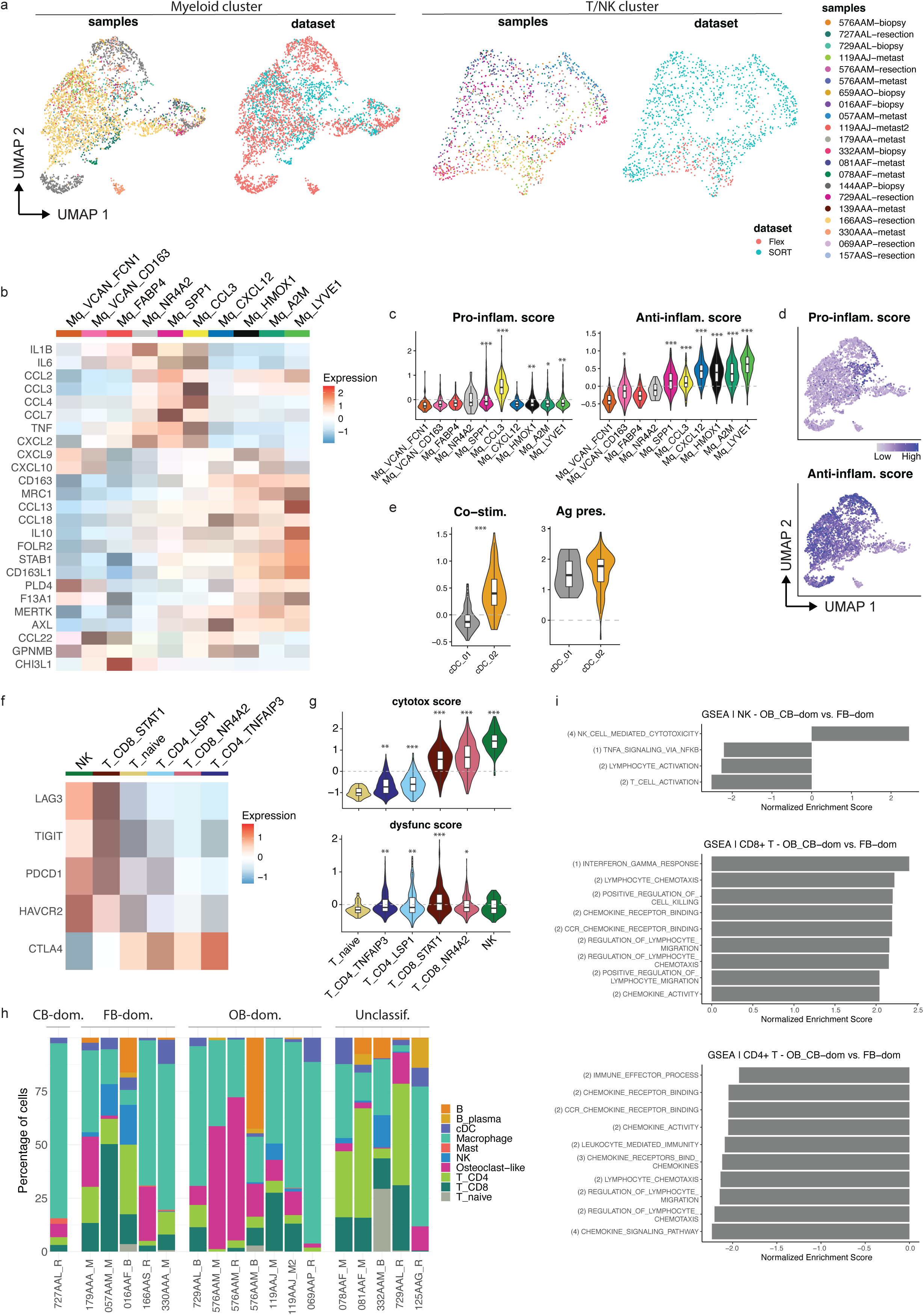
Immune cell subset characterization and functional states in OS. **a.** UMAP plot of myeloid cells (left) and T and NK cells (right), colored by sample ID and dataset. **b.** Heatmap showing the average expression of pro- and anti-inflammatory genes across myeloid cell subsets. **c.** Module scores for the pro- and anti-inflammatory gene sets shown in panel b. **d.** UMAP plots of the myeloid cluster, showing pro- and anti-inflammatory module scores. **e.** Module score of co-stimulatory and antigen presentation genes sets across cDC subsets. **f.** Heatmap showing average expression of inhibitory receptors across T and NK cell subsets. *\*Significantly upregulated with p<0.0001 using the Wilcoxon Sum Rank test with Bonferroni correction*. **g.** Dysfunction scores constructed using genes from panel f. Cell types are ordered from high to low score. *\*P<0.05, **P<0.01 ***P<0.001 versus T_naive*. **h.** Proportion of immune cell types per sample, grouped by dominant molecular tumor cell state. **i.** GSEA of NK cells and T cell subsets from OB_CB-dominant vs. FB-dominant samples. (1) indicates HALLMARK genesets, (2) indicates GO genesets, (3) indicates REACTOME genesets, and (4) indicates KEGG gene sets. Sample labels indicate biopsy (B), resection (R), or metastasis (M).

**Figure S7.**
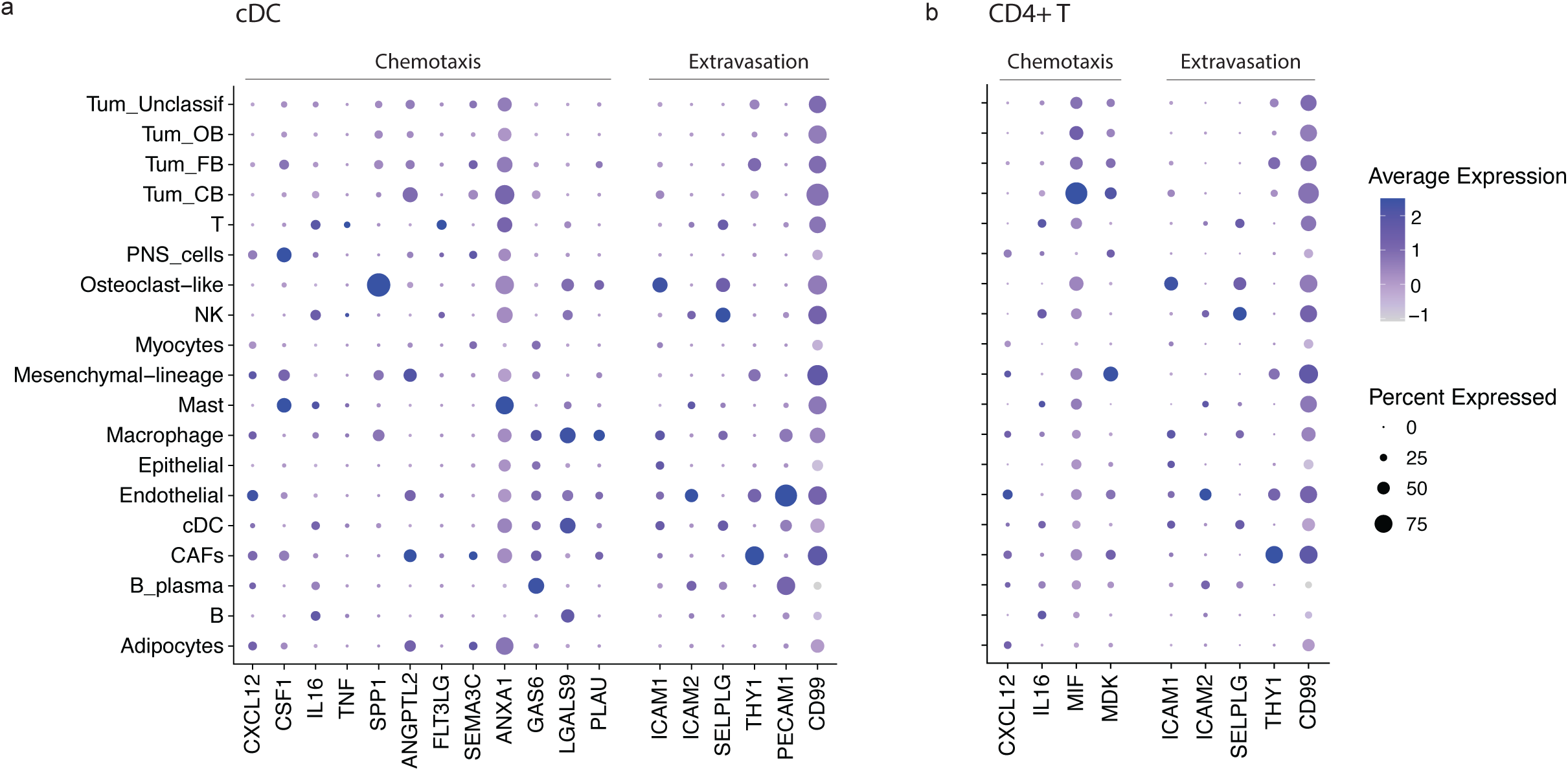
Expression of ligand-receptor interaction components across cell types. **a.** Dot plot showing expression of genes involved in ligand-receptor interactions targeting CD4+ T cells (as shown in Fig. 5d), by cell type. **b.** Dot plot showing the expression of genes involved in ligand-receptor interactions targeting cDCs (as shown in Fig. 5e), by cell type.

## Supplementary information

**Supplementary Table S1: Patients and samples characteristics**

Summary of tumor characteristics (primary site, metastasis, location, outcome), and sample preparation details (platform, number of cells per sample, and analysis).

**Supplementary Table S2: Ligand-receptor interactions of CD4+ T cells and cDCs with other cell types in FB-dominant tumors**

**Supplementary Data: Quantification of IHC staining**

Quantification of IHC staining for IBSP, ACAN, B7-H3, CSPG4, LRRC15, and ROBO1 across two OS tissue microarrays, showing percentage of positive cells and staining score for each marker.

